# Nucleolar stress controls mutant Huntingtin toxicity and monitors Huntington’s disease progression

**DOI:** 10.1101/2021.07.09.451766

**Authors:** Aynur Sönmez, Rasem Mustafa, Salome T. Ryll, Francesca Tuorto, Ludivine Wacheul, Donatella Ponti, Christian Litke, Tanja Hering, Kerstin Kojer, Jenniver Koch, Claudia Pitzer, Joachim Kirsch, Andreas Neueder, Grzegorz Kreiner, Denis L.J. Lafontaine, Michael Orth, Birgit Liss, Rosanna Parlato

**Author notes:** Correspondence to: Dr. Rosanna Parlato. shared last author (Michael Orth, Birgit Liss, Rosanna Parlato). **Abbreviations** CNN: convolutional neural network, DAPI (4′,6-diamidino-2-phenylindole), DBS: disease burden score, D1R: dopamine receptor 1, D2R: dopamine receptor 2, DTT: Dithiothreitol, ECL: Electrochemiluminescence, EDTA: Ethylenediaminetetraacetic acid, GAPDH: Glyceraldehyde 3-phosphate dehydrogenase, HD: Huntington’s disease; HPRT: Hypoxanthine-guanine phosphoribosyltransferase, ISH: *in situ* hybridization, Metap1: Methionine aminopeptidase 1, mRNA: messenger RNA, mHTT: mutant Huntingtin, NCL: nucleolin, NPM1: nucleophosmin 1, NSS: normal swine serum, PBS: phosphate-buffered saline, PFA: paraformaldehyde, PMSF: phenylmethylsulfonyl fluoride, PVDF: polyvinylidene fluoride, pre-rRNA: precursor rRNA, qRT-PCR: quantitative real time PCR, rRNA: ribosomal RNA, ROI: region of interest; STED: STimulated Emission Depletion, SUnSET: SUrface SEnsing of Translation, TBS: Tris-buffered saline, TFC: total functional capacity, TIF-IA: transcription initiation factor IA, TMS: total motor score, UHDRS: united Huntington’s disease rating scale.

## Abstract

Transcriptional and cellular stress surveillance deficits are hallmarks of Huntington’s disease (HD), a fatal autosomal dominant neurodegenerative disorder, caused by a pathological expansion of CAG repeats in the Huntingtin (*HTT)* gene. The nucleolus, a dynamic nuclear biomolecular condensate and the site of ribosomal RNA (rRNA) transcription, is implicated in the cellular stress response and in protein quality control. While the exact pathomechanisms of HD remain unclear, the impact of nucleolar dysfunction on HD pathophysiology *in vivo* is elusive. Here we identified aberrant maturation of rRNA and decreased translational rate in association with human mutant Huntingtin (mHTT) expression. Genetic disruption of nucleolar integrity in vulnerable striatal neurons of the R6/2 HD mouse model decreases mHTT disperse state in the nucleus, exacerbating the motor deficits. The protein nucleophosmin 1 (NPM1), important for nucleolar integrity and rRNA maturation, loses its nucleolar localization. NPM1 de-localization occurs in the striatum and in the skeletal muscle of the progressive zQ175 knock-in HD mouse model, mimicking the phenotype of HD patients in skeletal muscle biopsies. Taken together, we showed that nucleolar integrity regulates the formation of mHTT inclusions *in vivo*, and identified NPM1 as a novel, readily detectable peripheral histopathological marker of HD progression.

## Introduction

Dysregulation of rRNA biogenesis represents an emerging mechanism in several progressive neurodegenerative diseases characterized by proteinopathy (Aladesuyi Arogundade O *et al*., 2019; Amer-Sarsour F and Ashkenazi A, 2019; Herrmann D and Parlato R, 2018; Lee J *et al*., 2014; Parlato R and Bierhoff H, 2015; Schludi MH *et al*., 2015). Ribosomal RNA synthesis in the nucleolus - the most prominent nuclear compartment and a multilayered bio-molecular condensate - is tightly linked to the cell wellbeing, and it is highly responsive to cellular stress (Boulon S *et al*., 2010; Sharifi S and Bierhoff H, 2018). Nucleolar stress is a p53-dependent anti-tumoral surveillance pathway activated upon ribosome biogenesis dysfunction (Nicolas E *et al*., 2016). The shape of the nucleolus, its size, and the number of nucleoli per cell nucleus may change upon stress and in disease, reflecting changes in its function (Lafontaine DLJ *et al*., 2020). These properties have started to be explored as disease biomarkers (Derenzini M *et al*., 2009; Stamatopoulou V *et al*., 2018).

Huntington’ disease (HD) is caused by the expansion of CAG repeats in the exon 1 of the huntingtin (*HTT*) gene (Group THsDCR, 1993). This autosomal dominant mutation results in an abnormal polyglutamine expansion in the Huntingtin protein with toxic effects (Caron NS *et al*., 2018). Typical clinical hallmarks include motor, cognitive and psychiatric symptoms (Ghosh R and Tabrizi SJ, 2018). Dopaminoceptive medium spiny neurons of the striatum, a component of the basal ganglia, are particularly vulnerable to neurodegeneration, along with reduced connectivity in regional and whole brain cortico-caudate networks that highly correlate with cognitive and motor deficits (McColgan P *et al*., 2015). Other non-neuronal features include metabolic and immune problems, malfunction of skeletal muscle, and body weight loss (Bates GP *et al*., 2015; Dayalu P and Albin RL, 2015; Rodinova M *et al*., 2019). The length of the expanded CAG tract in the mutant *HTT* gene partially accounts for the variability in the clinical HD onset (Consortium. GMoHsD, 2019; Ghosh R and Tabrizi SJ, 2018; Stuitje G *et al*., 2017). Multiple pathophysiological mechanisms may contribute to HD (Saudou F and Humbert S, 2016). Mutant HTT (mHTT) protein forms nuclear and cytoplasmic inclusions that interfere with almost all aspects of cell physiology, from nuclear transcription dysregulation to mitochondrial dysfunction, and compromised quality control mechanisms, among many others (Francelle L *et al*., 2017; Liot G *et al*., 2017; McColgan P and Tabrizi SJ, 2018; Rai SN *et al*., 2019).

Previous studies showed that mHTT interferes with rDNA transcription and with the integrity of the nucleolus (Hilditch-Maguire P *et al*., 2000; Jesse S *et al*., 2017; Kreiner G *et al*., 2013; Lee J *et al*., 2011; Lee J *et al*., 2014; Trettel F *et al*., 2000; Tsoi H and Chan HY, 2013). In the striatum of the R6/2 transgenic mice, the *de novo* transcription of rRNA is impaired (Kreiner G *et al*., 2013). Because of its rapid pathological progression (Mangiarini L *et al*., 1996), this HD mouse model is commonly used for preclinical studies.

Several mechanisms have been proposed to explain how mHTT affects rRNA transcription (Jesse S *et al*., 2017; Lee J *et al*., 2011; Tsoi H and Chan HY, 2013). mHTT protein acts on the acetyltransferase CBP (CREB-binding protein), required for the activity of the RNA polymerase I (Pol I) (Lee J *et al*., 2011; Lee J *et al*., 2014). Moreover, *mHTT* messenger RNAs down-regulate rRNA transcription by interacting with the nucleolar protein nucleolin (NCL), that plays multiple roles in rRNA synthesis, ribosome biogenesis and nucleolar structure maintenance (Cong R *et al*., 2012; Tsoi H and Chan HY, 2013; Tsoi H *et al*., 2012). PGC-1alpha (peroxisome proliferator-activated receptor gamma co-activator 1alpha), a master regulator of mitochondrial biogenesis, which is transcriptionally repressed by mHTT, also controls ribosomal DNA transcription in the nucleolus (Jesse S *et al*., 2017). Importantly, Brain Derived Neurotrophic Factor known to sustain striatal neuron survival and downregulated in HD (Xie Y *et al*., 2010; Zuccato C *et al*., 2001), stimulates the activity of the transcription initiation factor-IA (TIF-IA), essential for the recruitment of the RNA Pol I at the ribosomal promoters (Gomes C *et al*., 2011).

An interesting role of the nucleolus is its involvement in protein quality control to prevent the irreversible aggregation of misfolded proteins, a mechanism often altered in several aggregate-forming neurodegenerative diseases (Frottin F *et al*., 2019; Nollen EA *et al*., 2001). In particular, the nucleolar protein nucleophosmin-1 (NPM1) appears to have a chaperone-like function in shielding the surfaces of potentially toxic aggregates (Woerner AC, Frottin F Science 2016).

Despite the multiple relationships between transcriptional, metabolic and quality control functions of the nucleolus and HD pathophysiological, the impact of nucleolar dysregulation on HD progression *in vivo* has not been systematically addressed. It also remained unexplored whether different disease stages are associated with context-specific changes in nucleolar transcription and integrity. Ultimately, cytomorphological nucleolar alterations related to HD pathophysiology in peripheral tissues might represent novel metabolic markers to monitor disease progression and treatment response.

To gain insight into the mechanistic relationship between nucleolar dysfunction and disease progression, we investigated the consequences of nucleolar stress on mHTT inclusions and motor symptoms in R6/2 mice. We used a functional genomic approach to inhibit RNA Pol I transcriptional activity in vulnerable striatal neurons of these mice. We have previously shown that the conditional ablation of *Tif-Ia* in medium spiny neurons of the striatum that express the dopamine 1 receptor (D1R) mimics a condition of progressive nucleolar stress *in vivo* (Kreiner G *et al*., 2013). Given that TIF-IA is targeted by several kinase cascades and integrates multiple signaling pathways, it represents an ideal molecular target to regulate rDNA transcription in a cell-type specific fashion (Drygin D *et al*., 2010; Parlato R and Kreiner G, 2013). We found that rRNA processing deficits are associated with loss of NPM1 nucleolar localization in a relevant neuronal model expressing human mHTT, suggesting a novel nucleolar stress-dependent pathophysiological mechanism. Finally, we reported similar subcellular distribution of NPM1 in skeletal muscle of a gradually progressive mouse model of HD, and in biopsies of HD patients at pre- and early symptomatic stages, establishing this characteristic as a reliable disease read-out.

## Results

### Expression of the human Huntingtin polyQ111 mutation alters nucleophosmin 1 localization, pre-rRNA processing and translation

To investigate the functional impact of mHTT expression on ribosome biogenesis and function, we analyzed a mouse cell model of Huntington’s disease derived from embryonic striatum and expressing a chimeric human-mouse mutant huntingtin (Trettel F *et al*., 2000). The StHdhQ111/Q111 cells (abbreviated “Q111/111”) contain the polyQ111 mutation encoded by the CAG expansion on both alleles in the Huntingtin gene. As control, we used Q7/Q7 cells that express a non-pathological 7-glutamine wild-type protein on both alleles **(Fig. 1)**. In this model, the localization of the scaffold protein nucleophosmin (NPM1) present in the nucleolar granular component, where late steps of ribosomal subunit assembly occur, was reduced (ca. 30%) in the Q111/111 cells in comparison with the control **(Fig. 1A, B)**. Importantly, nucleolin (NCL), another nucleolar protein, did not show any significance differences in abundance or distribution **(Fig. 1A, B)**.

**Figure 1:**
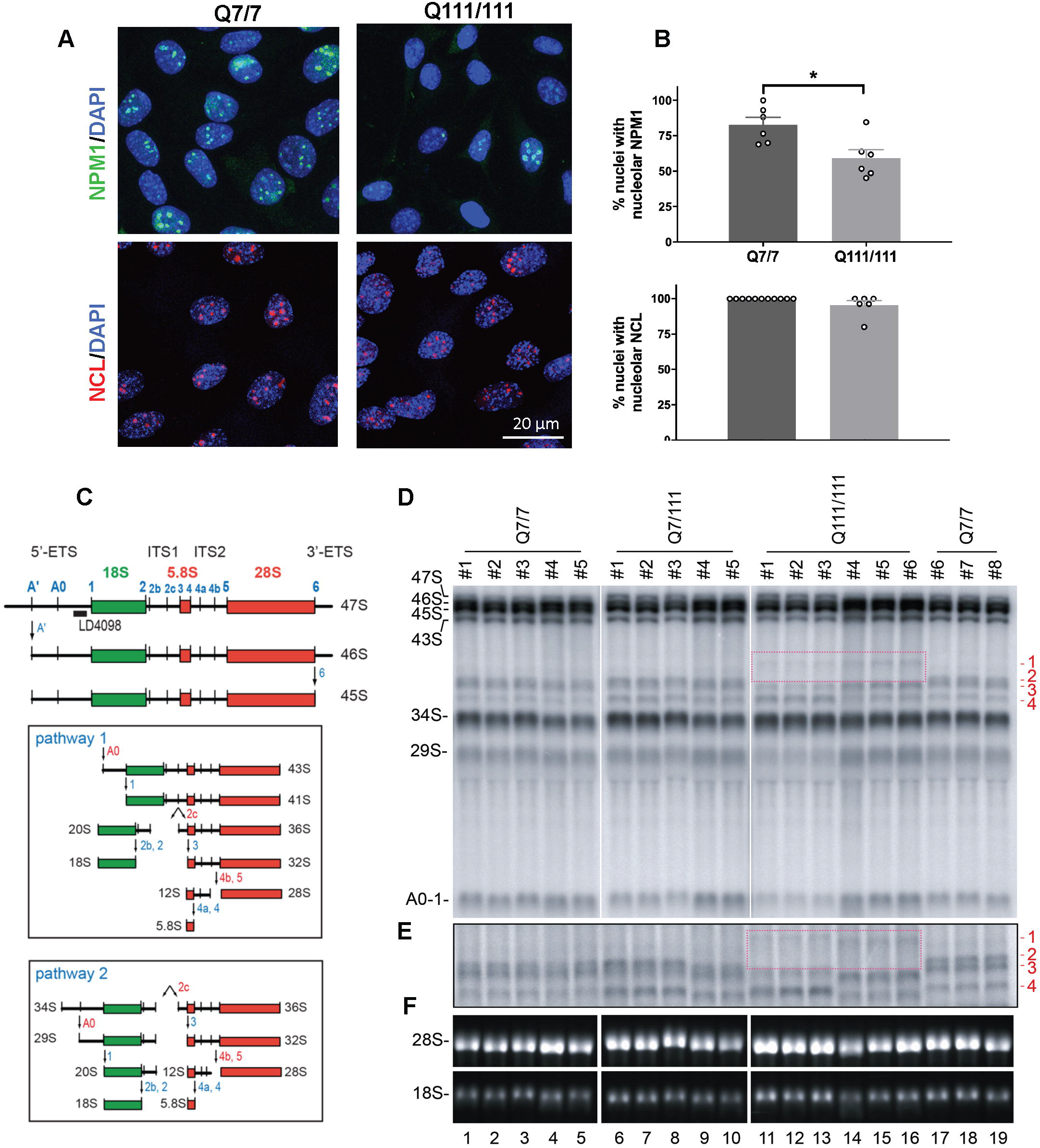
Cells expressing Huntingtin mutations display altered distribution of nucleolar NPM1 and alterations of pre-rRNA processing. (A) Representative confocal images of NPM1 and NCL immunofluorescence staining in Q7/7 and Q111/111 cells. (B) Quantification of the percentage of nuclei with nucleolar ring-like NCL and NPM1 signals. The percentage of nuclei with nucleolar NPM1 is significantly reduced in the Q111/111 cells by Mann-Whitney U test (p=0.014) (n: number of nuclei = 421 in Q7/7 and 634 in Q111/111; N: fields of view in two independent experiments = 6 for Q7/7 and Q111/111). Values represent mean values. Error bars represent SEM. * p<0.05. Detailed statistical information is included in the Supplementary Statistical Information file. (C) Mouse pre-rRNA processing pathway. Three out of four mature rRNAs (the 18S, 5.8S, and 28S) are encoded in a single polycistronic transcript synthesized by RNA polymerase I, the 47S. Mature rRNAs are embedded in 5’ and 3’ external transcribe spacers (5’- and 3’-ETS) and internal transcribed spacers 1 and 2 (ITS1 and 2) and are produced by extensive processing of the 47S. Processing sites (A’, A0, 1 etc.) are indicated in blue. There are two alternative processing pathways in mouse (pathways 1 and 2, boxed) according to initial processing in 5’-ETS or ITS1, respectively. (D) Total RNA was extracted, separated on denaturing high-resolution agarose gel, stained with ethidium bromide to reveal large mature rRNAs, or processed for Northern blotting. Species labeled ‘1 to 4’ (in red) are extended forms of the 34S pre-rRNAs that were not previously described. Species ‘2, 3, and 4’ are detected in the control cells (Q7/7). Species ‘1’ is only detected in the mutant cells (Q111/Q111). Species ‘1’ is formed at the cost of species ‘2’ (the upper band of the doublet). The boxed area highlights the appearance of species ‘1’ in Q111/Q111 cells, which is concomitant with the disappearance of species ‘2’. (E) The samples described in panel (D) were run in a longer migration to separate more efficiently the doublet corresponding to bands ‘2 and 3’. (F) Ethidium bromide staining revealing that the steady state levels of 18S and 28S rRNA are not grossly affected.

To test if mHTT expression impacts pre-rRNA processing, total RNA was extracted from cells expressing each construct, and separated on denaturing agarose gels. Mature rRNAs were visualized by ethidium bromide staining **(Fig. 1F)** and quantified from electropherograms. Precursor rRNAs were detected with specific radio-actively labelled probes **(Fig 1C-E)**.

We observed no gross alteration in the steady-state accumulation of the large mature rRNAs, the 18S and 28S rRNAs **(Fig. 1F)**. The canonical pre-rRNA processing intermediates (29S, 34S, 43S, 45S, 46S, and 47S) detected with probe LD4098 also accumulated normally **(Fig. 1D)**. However, close inspection of the imaging plates reproducibly revealed additional low-abundant pre-rRNA intermediates, referred to as ‘1’, ‘2’, ‘3’, and ‘4’ (in red in **Fig. 1D, E)**. Note that species ‘2’ and ‘3’ appeared as a doublet, which was well-resolved on longer migration (**Fig. 1E**, for example lanes 17-19). Species ‘2’, ‘3’ and ‘4’ visible in the reference cells (Q7/7) corresponded to extended forms of the 34S pre-rRNA. Cells that expressed the Q111/Q111 mutation, showed a novel band, labeled as ‘1’ (lanes 11-16, highlighted in a red box). Interestingly, when species ‘1’ was detected, ‘2’ was not, pointing to a precursor-product relationship that was specific to the Q111/111 cells. Remarkably, the cryptic species ‘1’ was not observed in the heterozygous cells (Q7/Q111), indicating that sufficient amounts of Q111/111 huntingtin must be expressed for processing to be altered.

We conclude that Q111/111 cells display qualitative differences in pre-rRNA processing, i.e. alterations in RNA cleavage kinetics. To investigate whether the processing alterations caused by mHTT expression impacts translation, we performed polysomal analysis in the Q7/7 and Q111/111 cells **(Supplementary Fig. 1A, B)**. Velocity gradient centrifugation allows to separate and to quantify free ribosomal subunits (40S, 60S), monosomes (80S), and polysomes, and it is a good proxy of global protein synthesis. Polysomal profiles revealed a deficit (ca. 20%, p=0.04) in global translation, analyzed by the reduction of polysomal fraction in Q111/111 cells, while the number of ribosomal subunits was similar to that of Q7/7 cells **(Supplementary Fig. 1A, B)**. This effect on translation was confirmed by use of the SUrface SEnsing of Translation (SUnSET) assay which monitors translation rates by labeling nascent proteins with the amino-acid analog puromycin **(Supplementary Fig. 1C, D)**. A significant reduction (ca. 30%, p=0.009) was observed in Q111/11 cells by comparison to control Q7/7 cells **(Supplementary Fig. 1C, D)**.

In summary, expression of mHTT in a cellular model of Huntington’s disease was associated with the specific loss of NPM1 nucleolar localization, the activation of cryptic pre-rRNA processing, and reduced global translation.

### Nucleolar stress exacerbates mHTT neuropathology and motor behavior deficits in the R6/2 model of Huntington’s disease

To investigate the impact of nucleolar stress on HD neuropathology, we genetically inhibited nucleolar function in striatal medium spiny neurons of the R6/2 transgenic mouse model of HD **(Fig. 2 and Supplementary Fig. 2)**. In particular, using the Cre-loxP system we conditionally ablated the transcription factor TIF-IA, essential for the activity of the RNA Polymerase I, in dopamine-receptive neurons of R6/2 mice, and we generated double mutant R6/2; TIF-IA^D1Cre^ mice (abbreviated “dm”) **(Fig. 2)**. We have previously shown that lack of TIF-IA in striatal medium spiny neurons in the TIF-IA^D1Cre^ mice inhibited rRNA synthesis and relocated nucleolar proteins to the nucleoplasm, mimicking a condition of nucleolar stress in striatal neurons by 4 weeks of age (Kreiner G *et al*., 2013).

**Figure 2:**
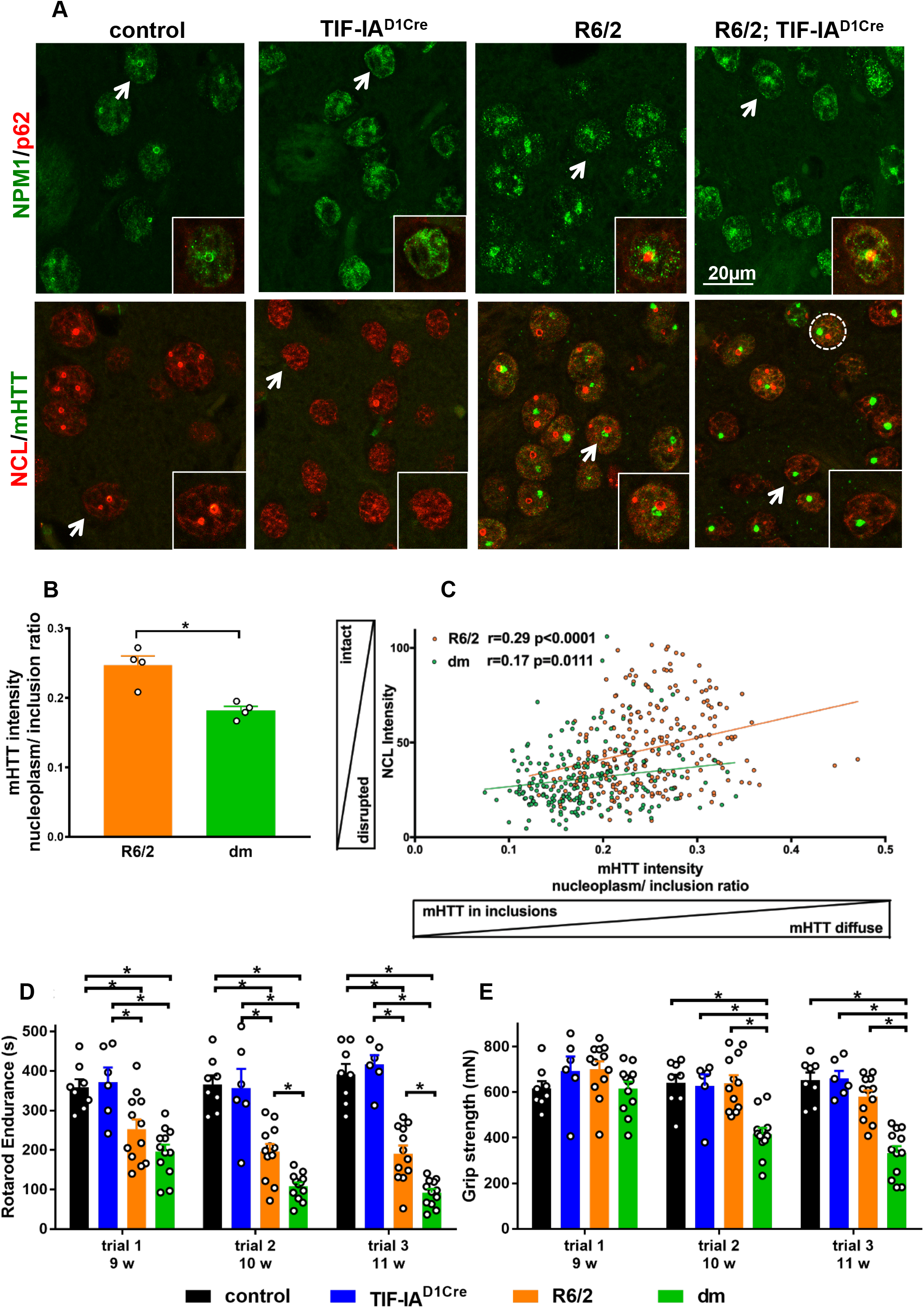
De-localization of NPM1 and NCL and altered nuclear distribution of mHTT inclusions upon irreversible induction of nucleolar stress in striatal neurons. (A) Representative confocal images of striatal sections from control, TIF-IA^D1Cre^, R6/2 and R6/2; TIF-IA^D1Cre^ double mutant (dm) mice at 9 weeks co-immunostained with antibodies against the nucleolar marker NPM1 and NCL, and p62 and mHTT (EM48), respectively. Insets are high magnification of the nuclei highlighted by the white arrow. Scale bar: 20 µm, 10 µm (insets). (B) mHTT nuclear distribution measured as the ratio between mHTT mean intensity in the nucleoplasm and in the nuclear inclusions in R6/2 (N=4) and dm (N=4) mice. mHTT intensity in the nucleoplasm of dm is significantly lower in comparison to R6/2 mice (Mann-Whitney U test, p = 0.03). (C) Higher nucleolar integrity assessed by NCL intensity in the nucleus correlates with a higher nucleoplasm/inclusion mHTT intensity ratio in R6/2 (n: number of nuclei=335, N: number of mice=4) and dm mice (n: number of nuclei=296, N: number of mice=4); (p<0.0001 for R6/2 and dm; Pearson coefficient r for R6/2 is 0.67 and for the dm 0.68). (D) Diagram showing endurance (in s) on an accelerating rotarod in the four experimental groups at different ages (9, 10 and 11 weeks, w). Significant variation between genotypes by two-way ANOVA (p<0.0001); significant differences between groups for each age were determined by Tukey’s multiple comparison: R6/2 vs. dm: p=0.012 at 10 weeks, p=0.003 at 11 weeks. N: number of mice, control: 8, TIF-IA^D1RCre^: 6, R6: 12, dm: 12. (E) Diagram showing the results of grip strength test in the four experimental groups at different ages (9, 10 and 11 weeks, w). Significant variation in age and genotype by two-way ANOVA (p< 0.0006); significant differences between groups for each age were determined by Tukey’s multiple comparison. N control: 8, TIF-IA^D1RCre^: 6, R6: 12, dm: 12. Legend: * p<0.05, ** p<0.01, *** p<0.001. Detailed statistical information is included in the Supplementary Statistical Information file.

Confocal microscopy on striatal sections immunolabeled by the nucleolar markers NPM1 and NCL showed that in control mice at 9 weeks, the nucleolus was visible as a circular structure within the nucleus **(Fig. 2A)**. At the same age in the TIF-IA^D1Cre^ mice, the percentage of DAPI labeled nuclei with NPM1 and NCL positive nucleoli was strongly decreased (ca. 90%) in comparison to control littermates **(Supplementary Fig. 2A).** In the R6/2 mice, both NPM1 and NCL maintained a ring-like organization, however NPM1 signal was diffuse **(Fig. 2A and Supplementary Fig. 2A)**. In addition, NPM1 surrounded the cargo protein p62/SQSTM1 immunopositive nuclear aggregates in both the R6/2 and R6/2; TIF-IA^D1Cre^ mice **(Fig. 2A)**.

Next, we compared the accumulation and distribution of mHTT in the R6/2 and in the R6/2; TIF-IA^D1Cre^ mice, by immunofluorescence using the EM48 antibody. As expected, mHTT was absent in striatal sections from control and TIF-IA^D1Cre^ mice **(Fig. 2A)**. In the R6/2 and R6/2; TIF-IA^D1Cre^ mice, mHTT was visible as intranuclear inclusions and scattered in the nucleoplasm **(Fig. 2A)**. The number and size of intra-nuclear inclusions containing mHTT was comparable between R6/2 and the R6/2; TIF-IA^D1Cre^ littermates **(Supplementary Fig. 2B)**. To gain further insight into the mHTT nuclear distribution upon induction of nucleolar stress, we measured the ratio between the intensity of mHTT signal in the nucleoplasm and in the nuclear inclusion bodies. Such ratio was about 30% lower in the R6/2; TIF-IA^D1Cre^ mice, suggesting that nucleolar stress alters mHTT distribution and reduces its diffuse state **(Fig. 2B)**. Disruption of nucleolar integrity determined by measuring nucleolin intensity in the DAPI stained nucleus in the R6/2 and in the R6/2; TIF-IA^D1Cre^ mice, correlated with lower intensity of the mHTT signal in the nucleoplasm **(Fig. 2C)**. This loss of diffuse nucleoplasmic mHTT was confirmed by an independent approach using immunohistochemistry with the EM48 antibody **(Supplementary Fig. 2C-E)**.

To further investigate whether nucleolar stress results in a more severe Huntington-like phenotype, we examined behavioral paradigms related to HD in control, TIF-IA^D1Cre^, in R6/2 and in TIF-IA^D1Cre^; R6/2 double mutant mice at 9, 10 and 11 weeks **(Fig. 2D,E)**. At 10 weeks, the double mutant mice performed significantly worse than the R6/2 mice, showing a more severe deficit in coordinating gait changes with increasing acceleration **(Fig. 2D)**. In line with these results, at 10 weeks the double mutants performed less well in the forelimbs grip strength test, in comparison to all other groups **(Fig. 2E)**.

These results indicated that NPM1 and NCL are differentially perturbed, and the pattern of mHTT accumulation points to a more advanced neuropathology in the double mutant mice.

### Reduced nucleolar localization of nucleophosmin 1 in the striatum of the late-onset zQ175 mouse model of Huntington’s disease

To monitor nucleolar activity and integrity in a gradually progressing model of Huntington’s disease, we analyzed heterozygous zQ175 knock-in mouse model of Huntington’s disease at pre-symptomatic (5 months) and symptomatic ages (10 month) (Carty N *et al*., 2015; Menalled LB *et al*., 2012). At 5 months, body weight and motor behavior of zQ175 heterozygous mice were comparable to that of control littermates **(Supplementary Fig. 3A-C**). In line with a more advanced disease stage, both dopamine D2-receptor *(D2r)* and D1-receptor (*D1r*) mRNAs were decreased at 10 months **(Fig. 3A)** (Menalled LB *et al*., 2012). Percentage of nuclei showing mHTT inclusions, area of the mHTT inclusions, and relative mHTT intensity in the nucleoplasm and in the inclusions confirmed the more advanced neuropathological stage at 10 months in comparison to 5 months in zQ175 mutant mice **(Fig. 3B-F** and (Carty N *et al*., 2015)). By qRT-PCR assays, RNA in situ hybridization (ISH) and Northern blots, we did not detect any significant differences in the 47S pre-rRNA, and mature 18S and 5.8S rRNA, indicating that rDNA transcription and mature rRNA *per se* are not impaired in the striatum of zQ175 mice at the two considered stages (**Fig. 4**).

**Figure 3:**
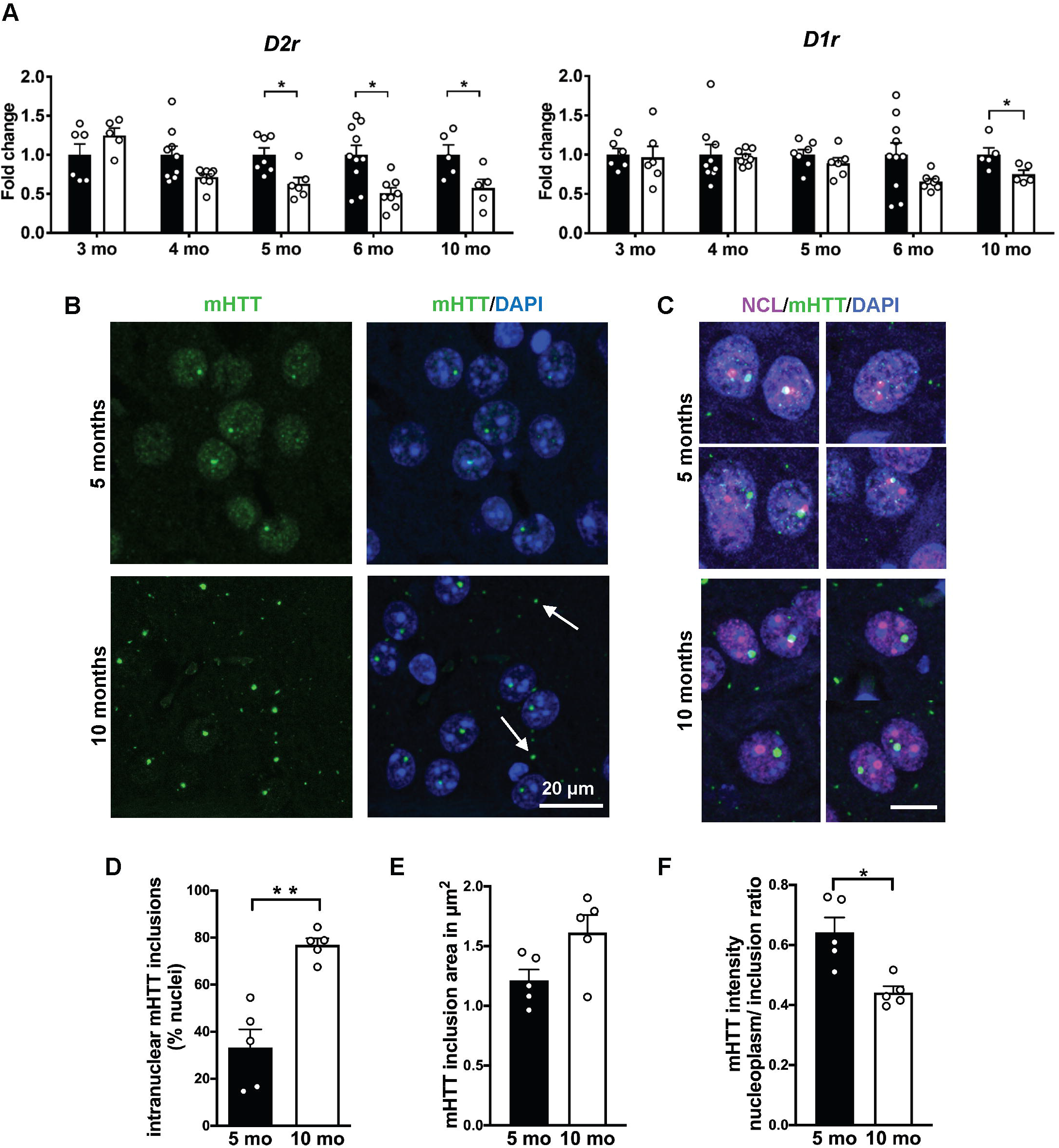
mHTT nuclear distribution changes at different stages in the striatum of zQ175 knock-in model. (A) Relative expression of *D2r* and *D1r* mRNA in the striatum by qRT-PCR at 3, 4, 5, 6 and 10 months (mo) in control (N=6, 8, 7, 10, 5) and zQ175 mutant (N=6, 8, 6, 8, 5) mice is expressed as fold change to respective controls. Significantly decreased relative expression of *D2r* at 5 (p= 0.014), 6 (p=0.012), 10 months (p=0.032) and of *D1r* at 10 months (p=0.032) by Mann-Whitney U test. (B, C) Representative confocal images of striatal sections from zQ175 mice at 5 and 10 months stained with antibodies against mHTT (EM48) and counterstained with DAPI to visualize the nuclei, and with antibodies against mHTT (EM48) and NCL. Scale bar: 20µm (B), 10µm (C). (D) Quantification of the nuclei with mHTT inclusion bodies in the zQ175 mice at 5 and 10 months shows a significant increase at 10 months by Mann-Whitney U test (p=0.008). (E) Non-significant statistical differences in the mean area of the mHTT inclusion signal by Mann-Whitney U test (p=0.09) between zQ175 mice at 5 and 10 months. (F) Nuclear distribution of mHTT is measured as the ratio between mHTT mean intensity in the nucleoplasm and in the nuclear inclusions at 5 and 10 months. mHTT intensity in the nucleoplasm at 10 months is significantly lower by Mann-Whitney U test (p=0.016). N: number of mice = 5; values represent mean values. Error bars represent SEM. * p<0.05, ** p<0.01.

**Figure 4:**
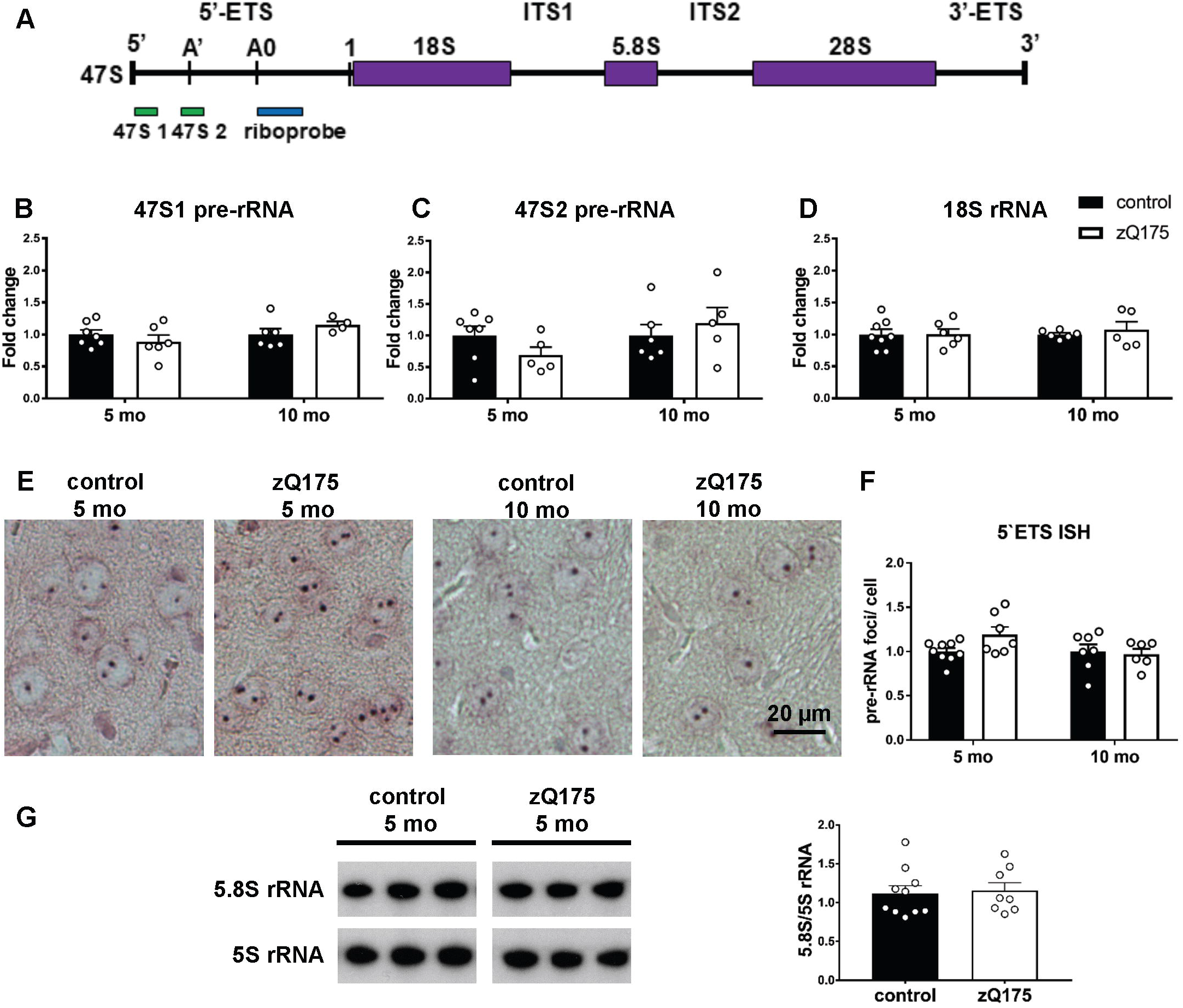
rDNA transcription is unaffected in striata of zQ175 mice at both pre- and early manifest stages. (A) Schematic representation of the 47S pre-rRNA transcript including the position of the primers used for qRT-PCR and of the riboprobe used for RNA ISH within the 5’-external transcribed spacer (5’-ETS). (B-D) Analysis of 47S pre-rRNA and mature 18S rRNA relative expression in the striatum by qRT-PCR at 5 and 10 months in controls (N= 6-8) and zQ175 (N= 4-6) shows no significant differences by Mann-Whitney U test. (E) Representative RNA ISH images of 5 and 10 month-old control and zQ175 mice showing the 47S pre-rRNA signals as blue punctuate signals within the nuclei in striatal sections. Scale bar: 20µm. (F) No significant difference in the average number of 47S pre-rRNA foci per striatal cell at 5 and 10 months in control (N= 9,7) and zQ175 mice (N= 7,6) by Mann-Whitney U test (p=0.09 at 5 months). (G) Representative Northern blots for the analysis of 5.8S rRNA in striata from controls (N= 10) and zQ175 (N= 8) at 5 months and densitometric analysis normalized by 5S rRNA. Error bars represent SEM.

The analysis of human BA4 cortex RNAseq data (GSE79666)(Lin L *et al*., 2016) and data from the striata and gastrocnemius of zQ175 mice at different ages (HDinHD.org)(Langfelder P *et al*., 2016) shows dysregulation of genes involved in rRNA transcription and pre-rRNA processing in this model at about 6 months **(Supplementary Fig. 3D,E)**. These observations along with the close interaction of mHTT with the nucleolus identified by immunofluorescence co-localization of NCL with mHTT in the zQ175 model **(Fig. 3C)**, led us to investigate changes in NPM1 nucleolar protein re-distribution **(Fig. 5)**. We analyzed the nucleolar distribution of NCL and NPM1 proteins by confocal microscopy, in the striatum of zQ175 mice at 5 and 10 months. The number of DAPI positive nuclei showing a distinct NPM1 nucleolar staining was significantly reduced (ca. 40% less, p=0.02) in striatal neurons of zQ175 mice at 5 months **(Fig. 5A, C)**. At 10 months, a ca. 20% reduction was observed, however it was not statistically significant (p= 0.15) **(Fig. 5A, C)**. Similar to the Q111/111 cells and R6/2 mice, we found no evidence for change of NCL in the zQ175 mice **(Fig. 5B, C)**. To further investigate the spatial distribution of NPM1 with respect to the mHTT nuclear foci, we performed super resolution microscopy in the striatum of zQ175 mice (STED, **Fig. 5D)**. We observed a close proximity between diffuse nucleoplasmic NPM1 and mHTT that appear intermingled at 5 months **(Fig. 5D, upper panels)**, while at 10 months, mHTT was mostly in the nuclear inclusion body **(Fig. 5D, lower panels)**, showing that re-distribution of nucleolar NPM1 precedes the formation of irreversible mHTT aggregates.

**Figure 5:**
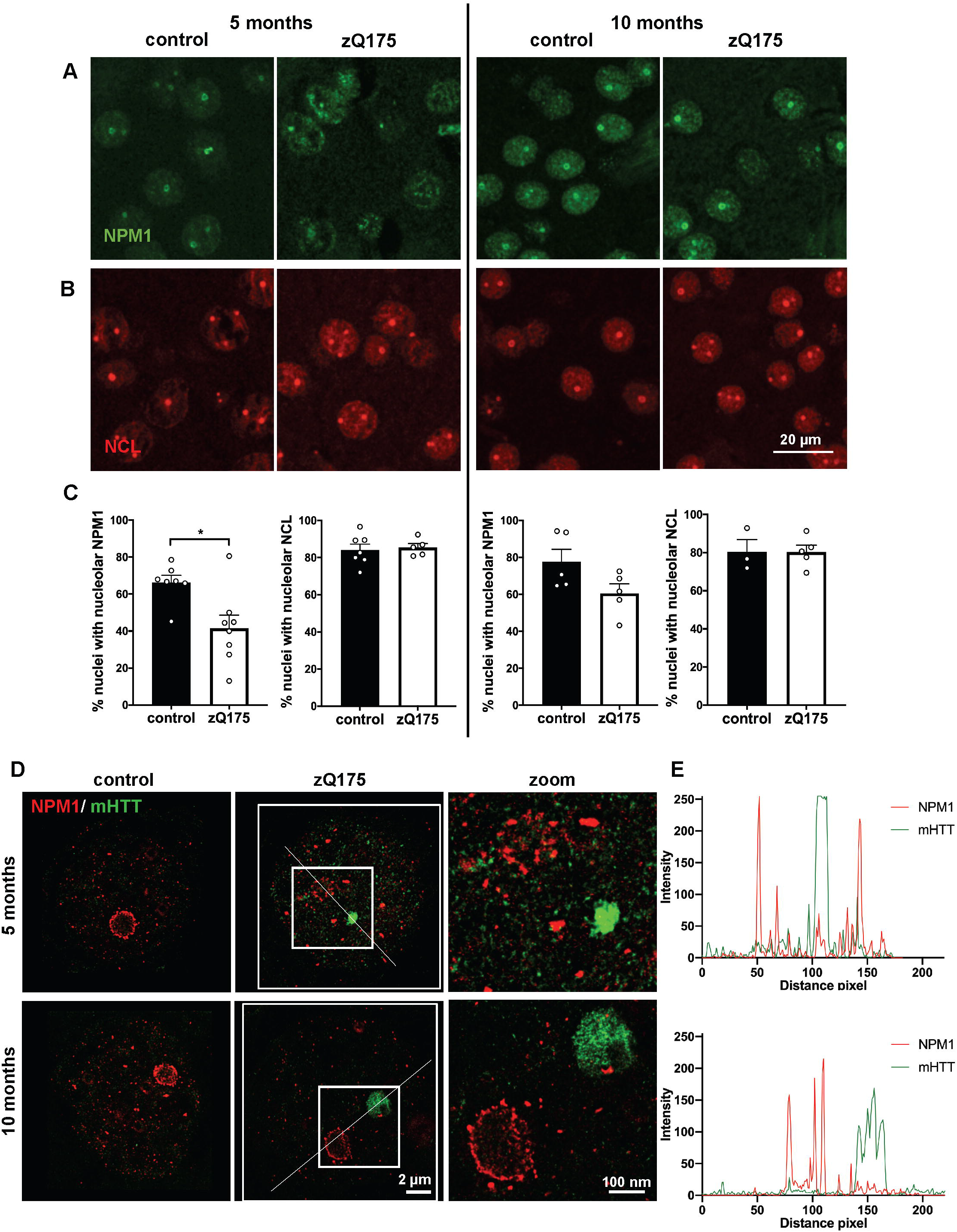
Redistribution of NPM1 in the striatum of pre-symptomatic zQ175 mice. (A,B) Representative confocal images of striatal sections stained for NPM1 (green) and NCL (red) in control and zQ175 mice at 5 and 10 months. Scale bar: 20 µm. (C) Quantification of the percentage of nuclei with nucleolar localization of NCL or NPM1 expressed as percentage of nuclei showing them as a circular nucleolar signal at 5 and 10 months in control and zQ175 mice (N: number of mice, control: 7,7,5,3 and zQ175: 8,5,5,5). Significant decrease in the number of nuclei showing nucleolar NPM1 in the zQ175 mice at 5 months in comparison to their respective controls by Mann-Whitney U test (p= 0.02). Values represent mean. Error bars represent SEM. * p<0.05. (D) Super-resolution (STED) microscopic images showing loss of NPM1 (red) ring-like organization and its association with disperse mHTT (green) signals at 5 months, but not at 10 months. (E) Line scans through the boxed regions containing the nucleolus and mHTT inclusion describe the distribution of NPM1 and mHTT signals in the zQ175 mice at both ages. A close proximity of mHTT and NPM1 signals at 5 months can be observed. Scale bar: 2 µm, zoom: 100 nm.

In summary, our results from striatal neurons in different models of Huntington’s disease identify nucleolar stress as a mechanism exacerbating disease progression, and NPM1 re-distribution as a marker for disease progression and mHTT aggregation.

### Reduced nucleolar localization of NPM1 in skeletal muscle of zQ175 mice

To explore the hypothesis that perturbed nucleolar homeostasis detected in the striatum by NPM1 nucleolar mislocalization represents a histopathological marker of Huntington’s disease progression, we analyzed skeletal muscle (quadriceps) of controls and zQ175 mice at 5 and 10 months **(Fig. 6A-C)**. At 5 months the number of DAPI positive nuclei showing either NPM1 or NCL signal was similar between zQ175 and control mice **(Fig. 6B)**. The area of the nuclei was also similar between control and zQ175 mutant mice at 5 and 10 months **(Supplementary Fig. 4A, B)**. However, in 10-month-old zQ175 mice, the nuclear area displaying NPM1 immunosignal was about 30% decreased **(Fig. 6C)**. Importantly, this reduction was specific, as it was not observed for NCL **(Fig. 6C)**. These findings suggest a nucleolar phenotype in skeletal muscle of zQ175 mice at 10 months. As a read-out for altered nucleolar function, we analyzed if rRNA synthesis was altered in skeletal muscle of zQ175 mice. We studied pre-rRNA transcription and mature 18S rRNA by qRT-PCR, and we detected at 10 months a significant (∼50%) decrease of 47S pre-rRNA in skeletal muscle of the zQ175 mice, in line with altered nucleolar function **(Fig. 6D)**.

**Figure 6:**
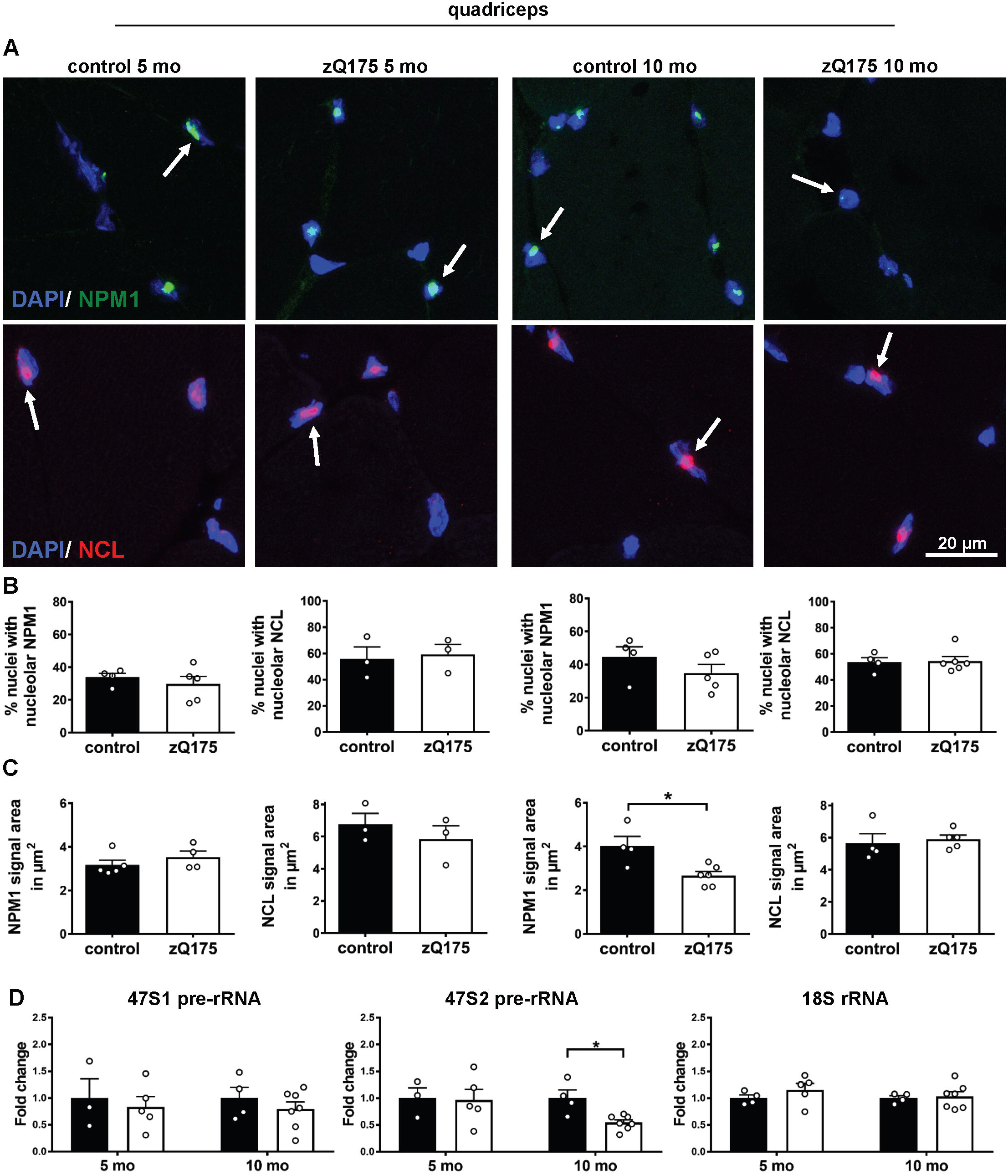
NPM1 nucleolar localization is altered in the skeletal muscle (quadriceps) of zQ175 mice at a symptomatic stage and downregulated rDNA transcription at an early manifest stage. (A) Representative confocal images of quadriceps cryosections stained for NPM1 (green) or NCL (red) in control and zQ175 mice at 5 and 10 months. Nuclei are labeled with DAPI (blue). The arrows point out to NPM1 or NCL signal. Scale bar: 20 µm. (B) Quantification of the percentage of nuclei with nucleolar localisation of NPM1 or NCL at 5 and 10 months (mo) in control (N, number of mice, NPM: 4, 4; NCL: 3, 4) and zQ175 (N: number of mice, NPM: 5, 5; NCL: 3, 6) mice shows no significant differences. (C) Quantification of the mean area of the NPM1 or NCL signal (in µm^2^) per DAPI positive nuclei at 5 and 10 months (mo) in control (N for NPM1: 5, 4; for NCL: 3, 4) and zQ175 (N for NPM1: 4, 6; for NCL: 3, 5) mice shows a significant reduction in nucleolar NPM1 signal at 10 months by Mann-Whitney U test (p= 0.038). * p<0.05. (D) Analysis of 47S pre-rRNA and mature 18S rRNA levels by qRT-PCR at 5 and 10 months in the skeletal muscle of controls (N= 3-4) and zQ175 (N= 5-7) expressed as fold change to respective controls. 47S pre-rRNA is significantly reduced using primer pair 47S2 on the zQ175 mice at 10 months by Mann-Whitney U test (p= 0.012). Values represent mean. Error bars represent SEM. * p<0.05.

Next, we asked whether the changes in NPM1 signal in skeletal muscle of zQ175 mice at 10 months were secondary due to a compromised function of striatal neurons, or whether they were associated with mHTT expression in the skeletal muscle. To this end, we analyzed skeletal quadriceps muscle from the TIF-IA^D1Cre^ mouse model, characterized by massive death of *D1r*-expressing medium spiny neurons at 3 months triggered by genetic induction of nucleolar stress in these neurons (Kreiner G *et al*., 2013). In this mHTT independent model of striatal neurodegeneration, nuclear NPM1 signals were not reduced but rather significantly elevated in skeletal muscle, compared to control littermate mice. These data suggest that the reduced NPM1 signal observed in the skeletal muscle of the zQ175 mice is linked to the peripheral effects of mHTT rather than to the degeneration of striatal neurons **(Supplementary Fig. 4D,E)**.

### Reduced nucleolar localization of NPM1 in skeletal muscle biopsies of Huntington’s disease patients

For the critical transition of our findings from Huntington’s disease mouse models to the human disease, we analyzed skeletal muscle biopsies from Huntington’s disease patients and control individuals. More precisely, we investigated the NPM1 and NCL fluorescence signal patterns in quadriceps muscle from biopsies of non-affected controls, pre-symptomatic and early-symptomatic Huntington’s patients **(Fig. 7A-C)**. Details about the gender, age at biopsy and length of the CAG repeat of the donors of muscle biopsies are provided in **Supplementary Table 1**. We analyzed the percentage of DAPI positive nuclei showing nucleolar NPM1 or NCL respectively, and the area of the NPM1 and NCL signal, similar as in the zQ175 mice. In addition, we established an automated approach to quantify NPM1 immunofluorescence signals, as a prerequisite for future systematic high-throughput histopathological assessments using NPM1 protein-distribution as a marker for disease progression.

**Figure 7:**
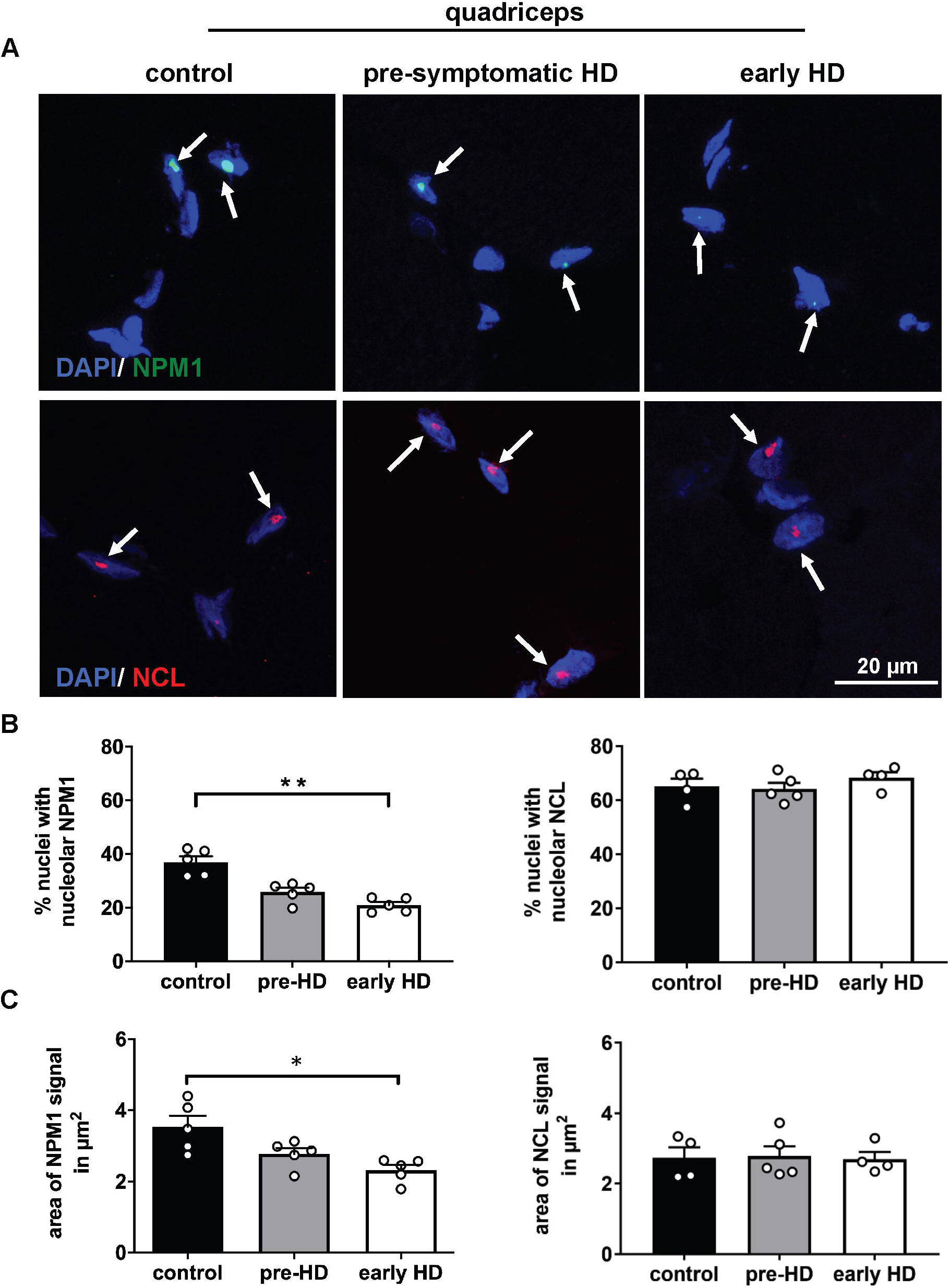
NPM1 signal is reduced in muscle (quadriceps) biopsies of early-Huntington’s disease patients. (A) Representative confocal images of human skeletal muscle (quadriceps) cryosections immunostained for NPM1 (green) or NCL (red) in Huntington’s disease patients at different stages and in age-matched controls. Nuclei are labelled with DAPI (blue). The arrows point to NPM1 or NCL signals. Scale bar: 20µm. (B) Quantification of the percentage of nuclei showing nucleolar NPM1 signal in Huntington’s disease patients in comparison with age-matched controls (N=5 for each group), and indicating significant decreased NPM1 in early Huntington’s disease (early-HD) by Kruskal-Wallis test and Dunn’s multiple comparison (p=0.003 early-HD vs. controls). No significant differences in the percentage of nuclei with nucleolar NCL. (C) Mean area of the NPM1 signal (in µm^2^) per DAPI positive nuclei in control, pre- and early-Huntington’s disease individuals (N=5 for each group). Statistically significant decrease of NPM1 signal area in early Huntington’s disease compared to controls by Kruskal-Wallis test and Dunn’s multiple comparison (p=0.01). No significant differences in the mean area of the NCL signal (in µm^2^) between the different groups. Error bars represent SEM. * p<0.05, ** p<0.01; detailed statistical information is included in the Supplementary Statistical Information file.

The images were first analyzed manually. Indeed, in early HD patients, the percentage of DAPI positive nuclei showing nucleolar NPM1, and also the area of the NPM1 signal were about 2- and 1.5-fold lower than those of controls, respectively **(Fig. 7B, C)**. In the pre-HD cohort, we observed a non-significant trend for reduced signals (p=0.14). A similar analysis performed for NCL showed no differences in these parameters at any stage (**Fig. 7B, C**). Nuclear areas were similar in all groups **(Supplementary Fig. 4C)**.

The same images were reanalyzed with an optimized automated approach based on the semantic convolutional neural network (CNN) learning, as indicated in the Materials and Method section. This analysis delivered similar results. Moreover, manually derived and automatically determined data displayed a strong correlation **(Supplementary Fig. 4F**). This automated approach represents a prerequisite for using NPM1 distribution in skeletal muscle as a marker for HD progression, and it opens the possibility to apply this analysis on a large scale of samples by digital pathology.

## Discussion

Nucleolar stress is associated with mHTT expression, and it is implicated in the response to cellular stress and in protein quality control. In this study, we provided the first *in vivo* evidences that this mechanism exacerbates motor phenotypes in a preclinical mouse model of HD, and that it alters mHTT intranuclear distribution. Aberrant rRNA processing in a striatal-derived cell model that expresses human mHTT indicated a novel mHTT-associated intrinsic dysfunctional mechanism. As summarized in the **Supplementary Table 2**, the nucleolar localization of the protein NPM1 was specifically reduced in the skeletal muscles of HD mice and in biopsies from HD patients, suggesting a novel candidate histopathological biomarker for monitoring HD progression in a peripheral tissue.

These findings are particularly important because changes in NPM1 distribution appear to be required for maintaining mHTT in a disperse state in the nucleoplasm. NPM1 and mHTT are known to interact (Culver BP *et al*., 2012; Shirasaki DI *et al*., 2012). NPM1 contains disordered and low-complexity sequences that may render this protein prone to interact with mHTT. Recent evidence showed that mHTT has distinct dynamic states in living cells, including fast diffusion rates, dynamic clustering and stable aggregation (Peskett TR *et al*., 2018). Studies in HEK293T cells transfected with artificial beta-sheet proteins, mimicking prefibrillar and fibrillar aggregate formation suggested that NPM1 might shield mHTT aggregate surface (Woerner et al, Science 2016). Our finding that the disruption of nucleolar integrity, and NPM1 release from the nucleolus, promote the loss of a diffuse mHTT state reconciles with the role of nucleolar integrity in protein quality control and formation of irreversible protein aggregates (Amer-Sarsour F and Ashkenazi A, 2019; Banski P *et al*., 2010; Frottin F *et al*., 2019; Nollen EA *et al*., 2001).

Previous studies showed that NPM1 is transiently upregulated in striatal neurons of the R6/2 transgenic mice before worsening of motor endpoints (Pfister JA and D’Mello SR, 2016), and that its nucleolar localization is partially lost in the same model at a pre-symptomatic stage (Kreiner G *et al*., 2013). Accordingly, the zQ175 mice show that NPM1 nucleolar localization is partially lost at a pre-symptomatic stage, further corroborating that changes in NPM1 subcellular distribution might be pathogenic. These findings are in agreement with the biphasic nature of nucleolar stress observed for example in amyotrophic lateral sclerosis (ALS). In this condition nucleolar stress occurs before the typical TDP-43 (TAR DNA-binding protein 43) proteinopathy (Aladesuyi Arogundade O *et al*., 2021).

In turn, mHTT interaction with NPM1 might exert toxicity by affecting the multiple functions of NPM1. On the other hand, other pathogenic effects can be hypothesized. NPM1 impairment leads to an accumulation of pre-rRNA and various processing intermediates, as shown by NPM1 depletion experiments in human cells (Tafforeau L *et al*., 2013). While NPM1 knockdown is toxic, its nuclear overexpression protects against mHTT-induced death in cultured cerebellar granule and cortical neurons (Pfister JA and D’Mello SR, 2016). We showed that the expression of human mHTT in striatal-derived cells is associated with qualitative alterations of ribosome production, namely the activation of cryptic processing sites, and that NPM1 loses its nucleolar distribution in the same model. At this stage, we cannot rule out that altered NPM1 affects pre-rRNA processing or sorting, as reported for another nucleolar protein fibrillarin in human cell cultures (Yao RW *et al*., 2019). Future studies should address whether such changes in the kinetics of RNA cleavage impact other facets of ribosome biogenesis, such as rRNA modifications or pre-ribosome assembly, and result in the production of ribosomes with altered translational abilities. Interestingly, in a genome-wide screen in yeast, expression of a mutant huntingtin fragment (Htt103Q) causes a dramatic reduction in expression of genes involved in rRNA metabolism and ribosome biogenesis (Tauber E *et al*., 2011). Moreover cells expressing HttQ138 show a decreased translation by the prion-like protein and translation regulator Orb2, sequestered by mHTT aggregates (Joag H *et al*., 2019). Recently, mHTT has been reported to suppress protein synthesis in the same HD cell model adopted here by a mechanism involving ribosome stalling (Eshraghi M *et al*., 2021). Increased translation has been however reported in a different progressive transgenic model of Huntington’s disease (Creus-Muncunill J *et al*., 2019). The phosphorylation of the eukaryotic translation initiation factor 4E (eIF4E) binding protein (4E-BP), an inhibitor of translation, increases in the striatum of R6/2 mice at a late manifest stage, supporting a time-specific dysregulation of translation (Creus-Muncunill J *et al*., 2019).

Our findings point not only to nucleolar stress as a contributor to Huntington’s disease, but also to re-distribution of nucleolar NPM1 as a cellular marker for Huntington’s disease progression. As summarized in **Supplementary Table 2**, there are tissue-specific differences within the same model (striatum vs. skeletal muscle), model-specific differences within the striatum (R6/2 vs. zQ175 mice) and age-specific differences within the muscle (zQ175 at 5 and 10 months) for nucleolar transcription and integrity. It remains to be analyzed whether these differences in nucleolar function and integrity correlate with different levels and / or state of mHTT (fibrils vs. inclusion bodies) and/or NPM1. These studies will be also important for the validation of NPM1 as a histopathological biomarker in longitudinal studies and for testing beneficial and adverse effects of therapeutic intervention on HD progression. In cancer research, changes in nucleolar size and shape are considered a reliable parameter to predict tumor growth and nucleolar proteins are evaluated as therapeutic targets (Derenzini M *et al*., 2000; Montanaro L *et al*., 2008; Nunez Villacis L *et al*., 2018). However, the value of the nucleolus as a potential biomarker is still neglected in the field of neurodegenerative diseases. We showed here, as a proof of principle, how the analysis of NPM1 could be automatized for clinical neuropathology applications, paving the way for high-throughput systematic analysis. A shared role for nucleolar stress in progressive neurodegenerative disorders, characterized by accumulation of intranuclear inclusions awaits future investigations (Amer-Sarsour F and Ashkenazi A, 2019; Latonen L, 2019; Lee KH *et al*., 2016; Rekulapally P and Suresh SN, 2019). The cell-specific and progressive modeling of nucleolar stress by using the TIF-IA conditional ablation is an important tool to investigate *in vivo* the early molecular, cellular, and behavioral responses to the loss of nucleolar homeostasis in genetic and pharmacological animal models (Domanskyi A *et al*., 2011; Evsyukov V *et al*., 2017; Kreiner G *et al*., 2013; Rieker C *et al*., 2011; Vashishta A *et al*., 2018).

In conclusion, we identified nucleolar stress as a disease mechanism contributing to HD pathogenesis, and in particular NPM1 distribution pattern in skeletal muscle as a novel promising candidate for developing an accessible and reliable biomarker for Huntington’s progression and possibly a target for therapeutics.

## Materials and Methods

### Statement regarding the ethical use of human material and animals

The ethics committee at Ulm University approved the study (Protocol number: 165/12), and written informed consent was obtained from each participant.

Procedures involving animal care and use were approved by the Committee on Animal Care and Use (Regierungspräsidium Karlsruhe, Germany, Animal Ethic Protocols: 34-9185.81/G-297/14 and 35-9185.81/G-102/16) in accordance with the local Animal Welfare Act and the European Communities Council Directives (2012/707/EU).

### Human skeletal muscle biopsies

Ten *HTT* CAG trinucleotide repeat expansion carriers and five healthy sex- and age-matched controls were recruited at the Department of Neurology of Ulm University (**Supplementary Table 1**). Participants had no contra-indications to muscle biopsy, e.g. a clotting disorder or abnormalities on electrocardiograms. *HTT* CAG repeat length was determined and participants were clinically assessed as described for the TrackOn and TRACK-HD studies (Orth M *et al*., 2017; Tabrizi SJ *et al*., 2009). The disease burden score (DBS) was calculated from each Huntington’s disease participant’s CAG repeat length and age according to the following formula: (CAG-35.5) × age (Penney JB, Jr. *et al*., 1997). Clinical assessment included the United Huntington’s Disease Rating Scale (UHDRS) motor part to derive the total motor score (TMS) (Group THsDCR, 1993) and the UHDRS total functional capacity scale (TFC). Potential Huntington’s disease participants were screened if they had a disease burden score of 250 or greater and either had no clinical signs of manifest motor Huntington’s disease (preHD) or were in TFC stages of 1 or 2 indicative of early motor manifest Huntington’s disease (earlyHD). **Supplementary Table 1** summarizes the participant demographic, number of CAG repeat, disease burden and UHDRS total motor score data.

Open biopsies of the *M. vastus lateralis* were obtained following local anesthesia. For immunofluorescence muscle was mounted on a piece of cork in TissueTek with fibers oriented perpendicular to the cork and then snap-frozen in liquid N_2_-cooled 2-methylbutane and stored at -80°C until sectioning.

### Mice

B6CBA-Tg(HDexon1)62Gpb/1J (R6/2) transgenic mice (CAG160) were imported from the Jackson Laboratories (Bar Harbour, ME). Htt^tmtm1Mfc^/190tChdi (zQ175 knock-in) mice (CAG198) were received from the CHDI Foundation by the Jackson Laboratories. For the experiments reported here, male and female mice were used, and wild-type and mutant littermates were analyzed. The zQ175 knock-in mice carry ca. 190 CAG repeats in a chimeric human/mouse exon 1 of the murine *Htt* gene (Menalled LB *et al*., 2012). The zQ175 mutation was kept in heterozygosity, to limit toxicity and mimic the most common genetic condition in Huntington’s disease (Menalled LB *et al*., 2012). The conditional knock-out of the nucleolar transcription factor *Tif-Ia* gene by the Cre-LoxP system in dopamine receptor 1 (D1R)-expressing cells (official nomenclature: B6.129.FVB/N-TIF-IA^tmGSc^Tg(D1RCre)^GSc^, abbreviated as TIF-IA^D1Cre^) was achieved as previously described (Kreiner G *et al*., 2013). Based on our breeding scheme crossing triple heterozygous mice with homozygous floxed *Tif-Ia* mice, we obtained 1 out of 8 R6/2 transgenic mice with the conditional loss of *Tif-Ia*. Mice were housed in a standard 12-h light/dark cycle and kept with ad libitum access to food and water. The analysis of the genotype was performed by PCR of tail snips as previously described (Kreiner G *et al*., 2013; Levine MS *et al*., 1999; Naranjo JR *et al*., 2016).

### Behavioral tests

Mice were tested for motor function on a rotarod apparatus (Ugo Basile, Gemonio, Italy) using a gradual acceleration protocol (4–40 rpm) up to maximal 8 min. Forelimb strength of all experimental groups was tested using a grip strength meter (Ugo Basile, Gemonio, Italy, model 47106), which automatically measured the force needed for the mouse to release its grip (Neureither F *et al*., 2017).

Rotarod and grip strength tests were performed 2 days after adapting to the respective tests at 9 weeks (trial 1), at 10 weeks (trial 2) and at 11 weeks (trial 3) for the experimental groups including control, TIF-IA^D1Cre^, R6/2 and their double mutant mice, and on the next day for the experimental groups including control and zQ175 mice (5 month-old). On the day of each trial each mouse was tested three times in intervals of 15 min. The mean endurance for the three runs (or recorded force, in millinewton) per mouse was calculated. The tests were performed blind for genotype.

### Tissue processing for RNA extraction and quantitative real-time PCR (qRT-PCR) in mice

Total RNA was isolated from dissected mouse striatum in the region comprised between Bregma 1.34 mm and -0.34 mm (Franklin P, 2008). Synthesis of complementary DNA (cDNA) with M-MLV Reverse Transcriptase (SuperScript III First Strand Synthesis Supermix, 18080-400, Thermo Scientific) was primed with random hexamers. For detection of pre-rRNA, either the first 130 nucleotides relative to the transcription start site (47S 1) were amplified using the 5’-ACTGACACGCTGTCCTTTCC and 5’-GACAGCTTCAGGCACCGCGA primers or a primer pair covering the first processing site (47S 2) was used: 5’-CGTGTAAGACATTCCTATCTCG and 5’-GCCCGCTGGCAGAACGAGAAG. To detect *Gapdh* mRNA (Glyceraldehyde 3-phosphate dehydrogenase) we used the following primers: 5’-CATGGCCTTCCGTGTTCCTA and 5’-GCGGCACGTCAGATCCA. Pre-rRNA and *Gapdh* mRNA amplification was performed using SYBR GREEN chemistry (SsoAdvanced Universal SYBR Green Supermix, 1725271, Bio-Rad), as previously described (Kiryk et al, 2013). The TaqMan gene expression assays (ThermoFischer Scientific) used for this study are reported in **Supplementary Table 3**. For each amplicon serial dilutions of cDNA were included in each run by a StepOnePlus instrument (Applied Biosystems) to generate standard curves that were used for relative expression quantification. Pre-rRNA expression in the striatum was normalized using the stably expressed reference gene *Gapdh* (glyceraldehyde-3-phosphate dehydrogenase), while for the TaqMan assays we used the stably expressed *Hprt* (hypoxanthine-guanine phosphoribosyl transferase) for striatal tissue and *Metap1* (methionine aminopeptidase 1) (Hering T *et al*., 2015; Hering T *et al*., 2017) for muscle tissue. Changes in relative expression were calculated as a fold change versus mean of respective control samples. All experiments were performed blind for genotype.

### Tissue processing for immunostaining in mouse and human tissues

Mice were sacrificed by cervical dislocation and brains and skeletal muscle (quadriceps) were immediately dissected. For immunofluorescence and immunohistochemistry, one brain hemisphere was fixed in 4% paraformaldehyde (PFA) in phosphate-buffered saline (PBS), pH7.4 overnight at 4 °C and paraffin embedded. Coronal sections (7 µm) were cut on a microtome (Leica 2235). The region of the striatum comprised between Bregma +0.74 mm and -0.34 mm was used for the histological analyses. Mouse skeletal muscle (quadriceps) was mounted on a piece of cork in Tissue Freezing Medium (12020108926, Leica) with fibers oriented perpendicular to the cork and then snap-frozen in liquid N_2_-cooled 2-methylbutane and stored at -80°C until sectioning. Sections were cut in transverse orientation to the fiber direction at 12 µm using a Leica CM1859 cryostat. Cryosections were fixed in 4% PFA for 15 min before staining. For antigen retrieval the slides were boiled in 1X citrate buffer (HK086-9K, Biogenex) for 3 min at 800 W in a microwave and then for 8 min at 400 W. Sections were blocked with 5% normal swine serum (NSS, S-4000, Vector) for 30 min. Primary antibodies were diluted in 5% NSS and incubated over night at 4°C in a humidified chamber. For immunohistochemistry, visualization of antigen-bound primary antibodies was carried out using a biotinylated secondary antibody together with the avidin-biotin system and the VECTOR peroxidase kit (PK-6100, Vector Laboratories) using both diaminobenzidine tablets (D4293, Sigma) as a substrate. Fluorescent-dye conjugated secondary antibodies were added and incubated for 30 min at room temperature. For double immunofluorescence staining, slides were incubated with a second primary antibody as described before. Nuclei were stained with DAPI (4′,6-diamidino-2-phenylindole) (1:10^6^, 62248, Invitrogen) for 10 min at room temperature. Slides were mounted using AquaPolymount (18606-20, Polysciences) and stored in the dark at 4°C until microscopic analyses. Primary antibodies for immunostaining were: anti-nucleolin (NCL, 1:500, ab70493, abcam) (Evsyukov V *et al*., 2017; Potapova TA *et al*., 2019), anti-nucleophosmin 1 (NPM1/B23, 1:100, MAB4500, Millipore) (Parlato R *et al*., 2006; Wang HF *et al*., 2011), sequestosome 1 (p62/SQSTM1) (1:100, P0067, Sigma), EM48 (1:100, MAB5374, Millipore) (Davies SW *et al*., 1997), S830 (1:1000; Neueder/Bates) (Moffitt H *et al*., 2009). For confocal microscopy, secondary antibodies Alexa 488 (A-21206, 1:100, Thermo Scientific) and Alexa 594 (A-21207, 1:100, Thermo Scientific) were used. For stimulated emission depletion (STED) microscopy, Alexa 594 (A-21207, 1:100 Thermo Scientific) and Star Red (STAR-RED-1056, 1:100 Abberior GmbH) were used.

### Mouse cell tissue culture conditions

The StHdh Q7/Q7, Q7/Q111, Q111/Q111 cells (striatal neuronal derived from Hdh7 wild-type and Hdh111 knock-in mice) (Trettel F *et al*., 2000) were cultured in Dulbecco’s modified Eagle’s medium medium (SIGMA) supplemented with Glutamax (GIBCO) and 10% inactivated fetal bovine serum (GIBCO) at 5% CO_2_ (Naranjo JR *et al*., 2016).

### Pre-rRNA processing analysis

5 µg total RNA from the StHdh Q7/Q7, Q7/Q111, Q111/Q111 cells (80% confluent grown in Petri dishes 6 mm diameter) was extracted, separated by 1.2% denaturing gel electrophoresis and processed for Northern blotting according to (Tafforeau L *et al*., 2013). The probe used was LD4098 (5’-ETS_A0-1, ACAATGACCACTGCTAGCCTCTTTCCCTT). Panels D and F were migrated at 60 V for 16 h; panel E, at 65 V for 24 h. The 28S/18S ratios were extracted from electropherograms (Agilent, RNA 6000 Nano Kit #5067-1511).

### Northern blot for 5.8S rRNA analysis

RNA was extracted from the entire striatum of one mouse brain hemisphere using the Trizol method (Invitrogen). 1-3µg were separated using 15% urea-PAGE (Invitrogen). Gels were stained with SYBR Gold (Invitrogen), and immediately imaged and transferred to Nytran SuperCharge membranes (Schleicher and Schuell), and UV crosslinked with an energy of 0.12 J. Membranes were hybridized in 5X SSC, 20mM Na_2_HPO_4_, pH7.4, 7% SDS, 1X Denhardt’s for at least 1 h at 42°C. For detection of 5.8S and 5S rRNA, membranes were incubated overnight (12–16 h) at 42°C with ^32^P-end-labelled oligonucleotides probes added in hybridization buffer with following sequence: 5.8S: 5’-GATGATCAATGTGTCCTGCAAT-3′; 5S: 5’-GGGTGGTATGGCCGTAGAC-3’. Following overnight incubation, the membranes were washed for 15 min at 43°C with 3X SSC, 5% SDS and for 15 min at room temperature with washing buffer (1X SSC, 1% SDS). Membranes were exposed on film, then stripped and re-hybridized with 5S probe for further analysis. Quantification of Northern blot signals from three independent experiments was performed on autoradiographs using Fiji/ImageJ (Schindelin J *et al*., 2012).

### RNA in situ hybridization for quantification of RNA foci

Non-radioactive RNA *in situ* hybridization (ISH) was performed on four to eight paraffin sections per mouse using a specific riboprobe hybridizing to regions in the leader sequence of the pre-rRNA as previously described (Rieker C *et al*., 2011). In brief after paraffin removal, sections were rehydrated and treated with proteinase K (10 µg/ml in 20 mM Tris/HCl, 1 mM EDTA, pH 7.2) for 7 min. Sections were incubated for 30 min in 2X SSC and in Tris/glycine buffer for at least 30 min until application of the hybridization mixture. The hybridization mixture was prepared as follows: 40% deionized formamide (Life Technologies, USA), 5X SSC, 1X Denhardt’s solution (Invitrogen, USA), 100 mg/ml salmon sperm DNA, 100 mg/ml yeast tRNA, 50 – 100 ng riboprobe, diluted in DEPC-H_2_O to a final volume of 100 µl per slide. Slides were incubated at 55 °C overnight. Post-hybridization steps were performed by washing the slides in 0.5X SSC/20% formamide at 60 °C and in NTE (0.5M NaCl, 10 mM Tris pH 7.0, 5 mM EDTA) at 37 °C. Slides were then incubated with 10 µg/ml RNase A in NTE for 30 min at 37 °C. Samples were washed in pre-warmed 0.5X SSC/20% formamide for 30 min at 60 °C and kept in 2X SSC for 30 min at room temperature. Sections were then placed in blocking solution (1 % blocking reagent, 11096176001, Roche, diluted in 100 mM maleic acid, 150 mM NaCl, pH 7.5) for 10 min at room temperature. Subsequently, sections were incubated with alkaline phosphatase-conjugated anti-digoxygenin antibody overnight at 4 °C. NBT/BCIP liquid substrate system for alkaline phosphatase (11681451001, Roche) was used for the staining reaction at room temperature in the dark. Incubation was stopped after 4 – 5 h. Sections were counterstained for 10 min with Nuclear Fast Red (Vector Laboratories, USA), and subsequently the coverslip was mounted with Kaiser’s Glycerol Gelatin (Merck, Germany). In situ hybridization and immunohistochemistry images were acquired by a Leica DM6B microscope with a DMC2900 camera using a 20X objective.

### Polysome profiling

5×10^6^ StHdh Q7/Q7 and Q111/Q111 cells were treated with cycloheximide (100 µg/ml) for 5 min and washed once in cold PBS/cycloheximide (100 µg/ml). The cells were lysed in 400µl lysis buffer (15 mM Tris–HCl, pH 7.4, 15 mM MgCl_2_, 300 mM NaCl, 1% Triton X-100, 100 µg/ml cycloheximide, 1X complete protease inhibitors (Roche), 0.1% β-mercaptoethanol). As RNAse inhibitor we used RNasin (200U/ml) (Thermo Fisher Scientific). The lysates were centrifuged at 10,000 g for 10 min at 4 °C, and the supernatants were applied to linear 17.5–50% sucrose gradients in 20 mM Tris–HCl (pH 7.4), 5 mM MgCl_2_, and 150 mM NaCl. Centrifugation was carried out at 36,000 rpm for 2.5 h at 4°C in a Beckmann SW60 rotor. Gradients were eluted with an ISCO UA-6 gradient fractionator, and polysome profiles were recorded by continuously monitoring the absorbance at 254 nm. The fraction of ribosomes engaged in translation was calculated by dividing the area under the polysomal part of the curve by the total area.

### SUrface SEnsing of Translation (SUnSET) assay

StHdh Q7/Q7 and Q111/Q111 cells (90% confluent grown in Petri dishes 6 cm diameter) The cells were incubated with 1µg/ml of puromycin for 30min and then analyzed to detect the puromycin-labeled peptides by immunoblotting. The puromycin concentration was determined by preliminary experiments of cell viability at 0.1-1.5 µg/ml. For immunoblotting total extracts were prepared by re-suspending cells in RIPA buffer (25 mM Tris-HCl pH 7.6, 150 mM NaCl, 5 mM EDTA (Ethylenediaminetetraacetic acid), 1% Triton, 1% sodium deoxycholate, 0.1% SDS), 1 mM PMSF (phenylmethylsulfonyl fluoride), 1 mM DTT (Dithiothreitol), 1X proteinase inhibitor (ROCHE complete, EDTA free). Protein quantification was performed by using the Thermo Scientific Pierce BCA protein Assay kit (detergent compatible formulation). The whole extracts (20 µg) were loaded on 10% SDS-PAGE gel and blotted on PVDF (Polyvinylidene fluoride) membrane (Biorad) with wet apparatus (Biorad), for 60 min at 400 mA. After blocking in 5% dry milk in TBS, the membrane was incubated overnight with the appropriate primary antibodies. The primary antibodies used were: anti-puromycin (Millipore MABE343, 1:1000) and anti-beta actin (sc-47778, Santa Cruz Biotechnology, 1:1000). The secondary antibody was anti-mouse HRP (horseradish peroxidase)-linked (NA931, Amersham ECL Mouse IgG, HRP, GE Healthcare Life Science) diluted 1:10000 in 5% dry milk in PBS, and incubated for 1h at room temperature. After ECL (Electrochemiluminescence) assay (Pierce ECL Western Blotting Substrate cat. 32106 Thermo Fisher Scientific), the membrane was incubated with the film (Kodak) for protein detection. As additional loading control, the same amount of total protein extracts were loaded on 10% SDS-PAGE and then stained with Coomassie Blue R-250 (Sigma). Protein expression was quantified by densitometric analysis with the ImageJ Software.

### Confocal and STED super-resolution microscopy

Confocal images of the striatum and muscle sections were obtained as z-stacks using a confocal scanning microscope (Leica TCS SP8) with a 63X/1.32NA oil objective and Leica LAS X imaging software. STED images were acquired as z-stacks in 0.5 µm steps at the Stedycon (Abberior Instruments GmbH) with a 100X/1.4NA oil objective. STED images were deconvoluted by the Huygens software (Scientific Volume Imaging, Hilversum, The Netherlands).

### Image analysis

All microscopic images were analyzed by an experimenter blinded to the genotype and age using the Fiji software (Schindelin J *et al*., 2012). The number and area of nucleolar markers were analyzed, as previously described (Evsyukov V *et al*., 2017; Kiryk A *et al*., 2013; Kreiner G *et al*., 2013; Tiku V *et al*., 2018). The percentage of nuclei showing NCL and NPM1 in the nucleoli was determined by counting the number of DAPI positive nuclei showing a circular nucleolar signal. The mean area of the nucleolar marker signal was determined by circling the signal area in each DAPI positive nucleus, considering in total only the double-labelled nuclei (Tiku V *et al*., 2018). The percentage of DAPI positive nuclei showing intranuclear mHTT inclusions and their mean area was measured in a similar way. For the semi-quantitative analysis of the signal mean intensities, nuclei labelled with DAPI were identified and marked in maximal projection images and the DAPI negative region outside was considered as the background. A cut-off of 50 µm^2^ was applied to exclude smaller fragmented DAPI positive signals. The DAPI positive area identified the region of interest (ROI). The mean intensities of NCL in the entire nucleus and the respective background were measured in each of the 10 planes of the z-stack, and the intensities of every plane were summed up. The intensity of the signal in the ROI was subtracted by the background intensity for each nucleus. The relative mHTT subcellular distribution was defined by the ratio between mHTT mean intensities in and outside the nucleus (Gasset-Rosa F *et al*., 2017). The intranuclear distribution was defined as the ratio between mHTT intensity in the nucleoplasm and in the inclusions (Frottin F *et al*., 2019). The region of the nucleus not occupied by the mHTT inclusions was considered as the nucleoplasm. For this analysis the mHTT inclusions were marked on the maximal z-projection. Intensities were measured across the whole z-stack and summed up. For each nucleus the intranuclear mHTT distribution was calculated by dividing the mHTT intensity in the nucleoplasm by the mHTT intensity in the inclusion.

For the quantification of the number of pre-rRNA foci by in situ hybridization the signals produced by labeling the riboprobe with digoxigenin were counted and normalized to the number of nuclei identified by counterstaining with the Nuclear Fast Red, as previously reported (Evsyukov V *et al*., 2017).

The manual segmentation of NPM1 positive nucleoli in human skeletal muscle biopsies was automated by a semantic image segmentation algorithm based on deep learning. The neural network used here is a fully convolutional neural network (FCN) using exclusively convolutional operations and it can thus be applied to images of arbitrary size in a computationally efficient way (Shelhamer E *et al*., 2017). The detailed network architecture used here was developed and provided by Wolution GmbH & Co. KG, through its scientific image analysis platform, and it uses five convolutional layers as well as activation functions of ReLu type. The power of such architectures for medical applications was previously demonstrated (Havaei M *et al*., 2017). The results of the percentage of NPM1 positive nuclei and of the NPM1 signal area achieved by the manual and automated approach were then correlated. Line profiles were measured after acquisition using ImageJ. A line through the nucleolus was drawn and measured using the Plot Profile function, as previously described (Riback JA *et al*., 2020; Taslimi A *et al*., 2014; Zhu L *et al*., 2019).

### Statistics

Statistical analysis was performed with Graphpad Prism 7.04 software. Mean values per sample were used for statistical analysis. Datasets were analyzed for their statistical significance using non-parametric unpaired, two-tailed Mann-Whitney U (MWU) test or using the Kruskal-Wallis test followed by Dunn’s post-hoc analysis for multiple comparisons (Altman DG *et al*., 1983). The MWU tests could not be applied for data shown in Fig. 4 due to no variation of the data points for the control group. Behavioral data were analyzed by two-way ANOVA with Tukey’s post-hoc analysis. For all tests statistical significance level was set at p<0.05. Complete details about statistical analysis are provided as **Supplementary Statistical Information**.

### Data availability

The raw data that support the findings of this study are available under “Supplementary statistical information” and from the corresponding author upon reasonable request.

## Supporting information

Suppl Figures with legends and tables

List of gene in Suppl Fig 3

Suppl stat information tables

## Acknowledgements

We thank all human participants for their participation and donation of tissue samples, and CHDI for providing the zQ175 mice. We also thank Dr. Gillian Bates for providing the S829 antibody, Dr. Jose Naranjo for providing the StHdh cells and protocols, Dr. Xunlei Zhou for her initial help with cell cultures. Barbara Kurpiers for her assistance with the behavioral tests, Dr. Nina Ullrich for assistance with the Sunset assay, and Dr. Rosario Piro for assistance with statistical analysis.

## Author contribution

AS, RM, STS, LW, FT, DP, CL, KK, JK, TH, CP, RP: performed experiments; AS, MO, RM, STS, FT, DP, CL, JK, CP, AN, GK, JK, DLJ, BL, RP: analyzed data; AS, MO, AN, JK, DLJ, BL, RP: supervised experimental work; AN, GK: contributed antibodies and mouse lines; MO: contributed human muscle biopsies and study design; MO, RP: conceived the study; AS, MO, DLJ, BL, RP: wrote the manuscript. The first draft of the manuscript was written by RP and all authors commented on previous versions of the manuscript. All authors read and approved the final manuscript.

