## Supplementary material for "Nucleolar stress controls mutant Huntingtin toxicity and monitors Huntington’s disease progression": Suppl Figures with legends and tables

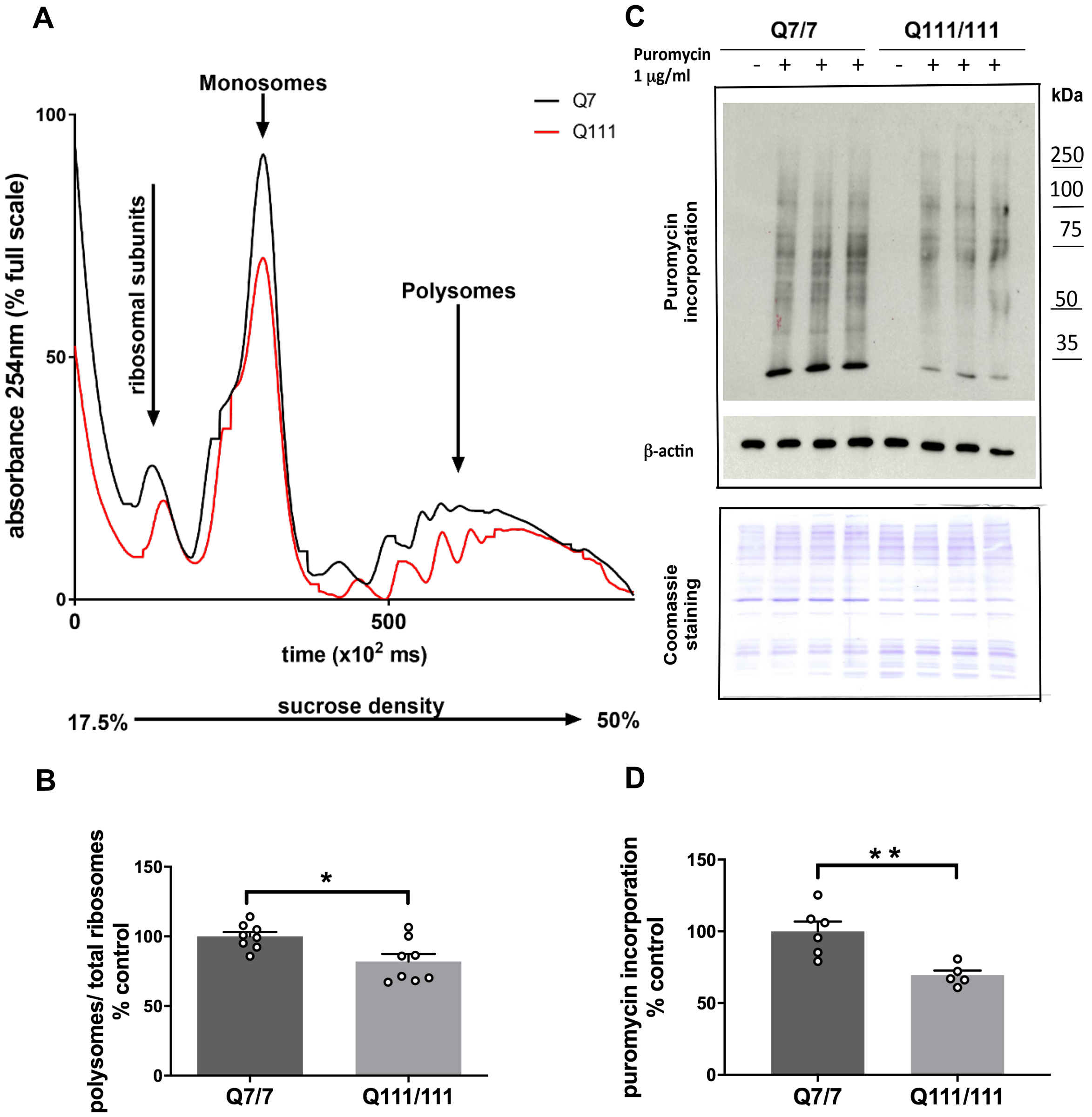


Suppl. Figure 1

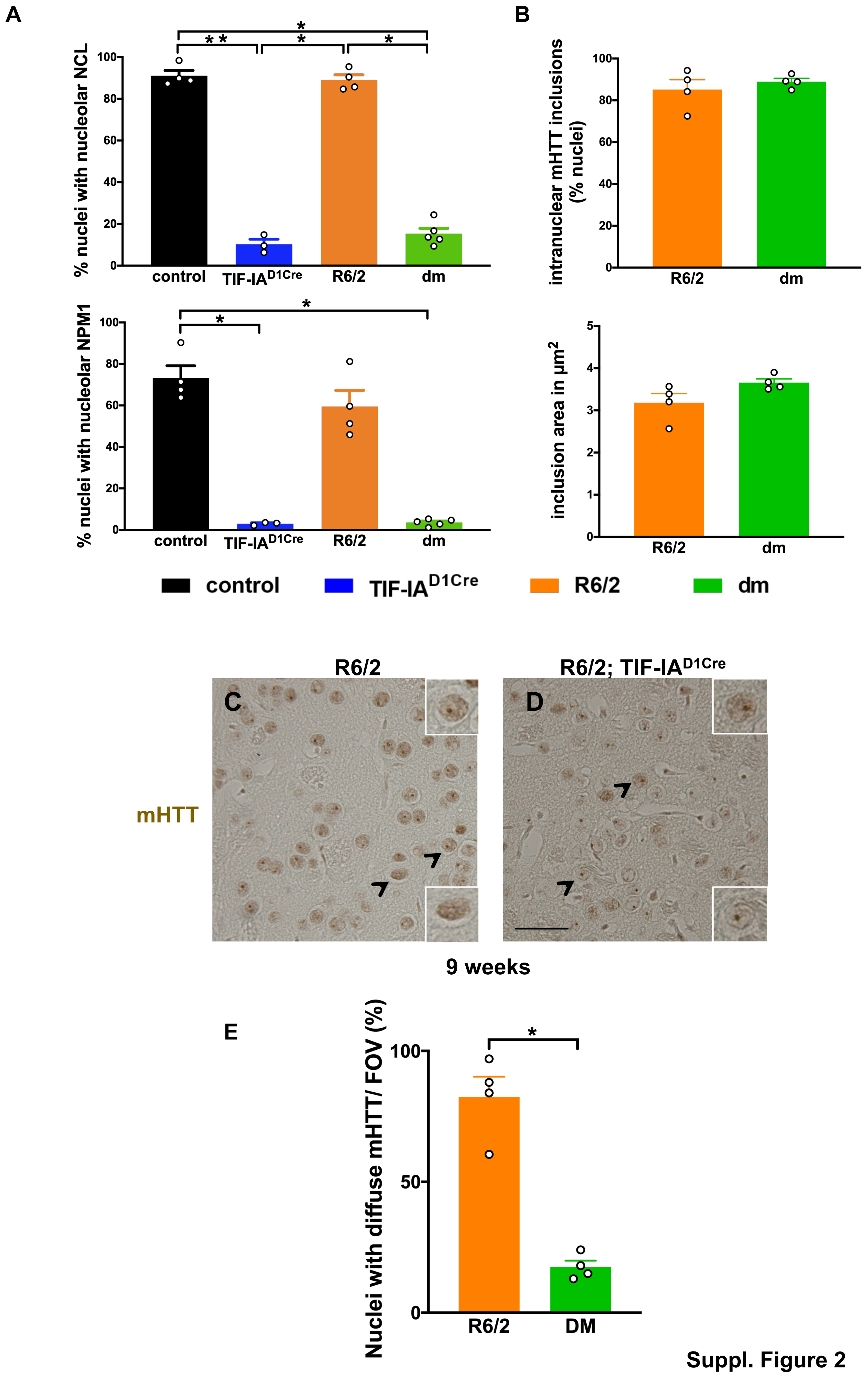


Suppl. Figure 2

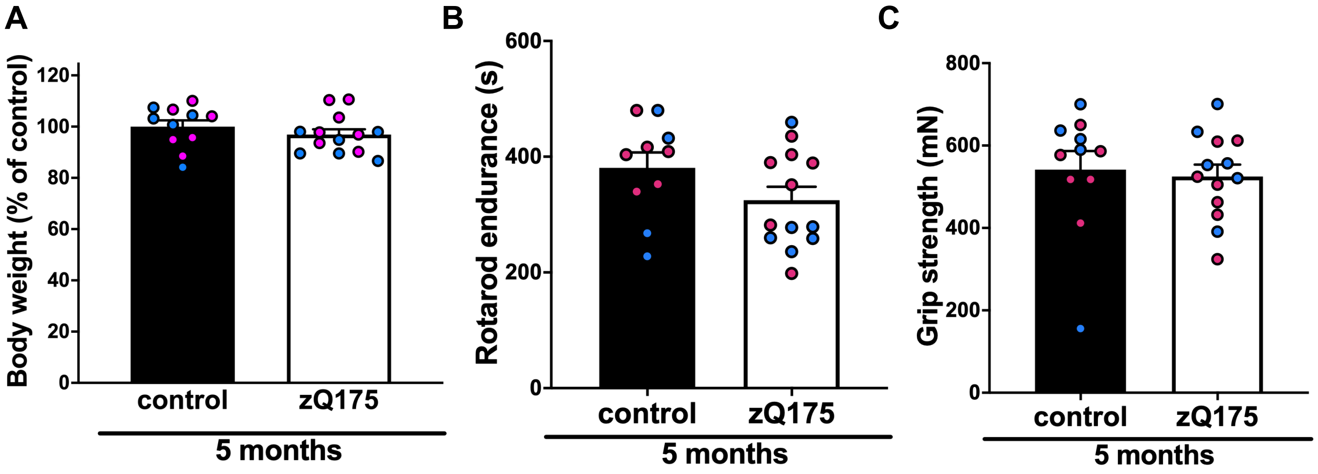


**D** rRNA transcription

| Gene | human BA4 cortex | zQ175 striatum 2 mth | zQ175 striatum 6 mth | zQ175 striatum 10 mth | zQ175 gastrocnemius 6 mth |
|---|---|---|---|---|---|
| *RRN3* | 0.29313315 | | | | |
| *UBTF* | | -0.1757289 | -0.320152 | -0.252921 | |
| *GTF2H2* | | | | -0.129181 | |
| *ERCC6* | | | -0.1514219 | -0.1519405 | |
| *TAF1C* | | | -0.0894431 | | |
| *GTF2H3* | | | -0.0754702 | | |
| *ERCC2* | | | 0.06336042 | | |
| *PCNA* | | | 0.09297059 | | |
| *KAT2B* | | | 0.10771751 | 0.12790748 | |
| *GTF2H5* | | | 0.10827059 | | |
| *DDX21* | | | 0.11621982 | 0.18567179 | |
| *TOP1* | | | 0.11746182 | | |
| *POLR1A* | | | 0.14489715 | | |
| *ERCC3* | | | 0.17364773 | 0.23119398 | |
| *MYO1C* | | | 0.18749787 | 0.2273397 | |
| *POLR1E* | | | 0.21095236 | | |
| *XRCC5* | | | 0.24006573 | 0.19054009 | |

**E** pre-rRNA processing

| Gene | human BA4 cortex | zQ175 striatum 2 mth | zQ175 striatum 6 mth | zQ175 striatum 10 mth | zQ175 gastrocnemius 6 mth |
|---|---|---|---|---|---|
| *NHP2* | | | -0.3335672 | -0.2729504 | |
| *RPL31* | | | -0.2410299 | -0.2047134 | |
| *LTN1* | | | -0.0901754 | | |
| *RPS19* | | | -0.0821946 | | |
| *SBDS* | | | -0.0782517 | | |
| *EXOSC10* | | | -0.0749136 | | |
| *RPS24* | | | -0.0661457 | | |
| *EFL1* | | | 0.08088634 | | |
| *RPL18* | | | 0.09356379 | | |
| *NPM1* | | | 0.09983137 | | |
| *TCOF1* | | | 0.1492631 | | |
| *DIS3* | | | 0.17113883 | 0.1884542 | |
| *FBL* | | | 0.22095978 | 0.20631843 | |

log2 fold change

< -1   0   > 1

**Suppl. Figure 3**

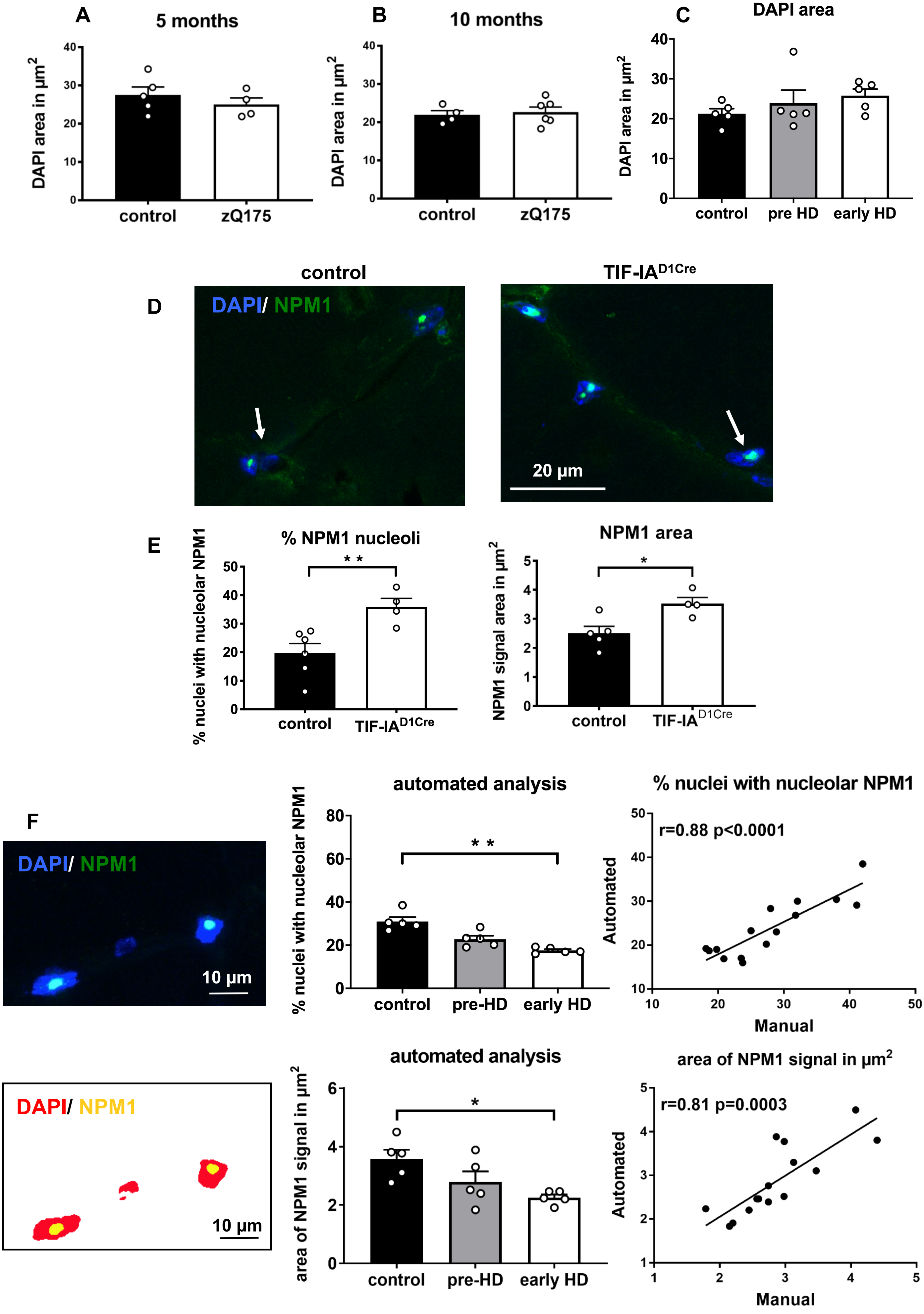


Suppl. Figure 4

### Supplementary Information

#### Supplementary Figure Legends

**Supplementary Figure 1: Expression of mutant Huntingtin in cells is associated with translational deficits.** (A) Representative polysome profiles of the Q7/7 and Q111/111 cells. (B) Ratio of polysomes to total ribosomes. Statistical significance by Mann-Whitney U test, * $p<0.05$. (C) Representative Western blot showing newly translated proteins in the Q7/7 and Q111/111 cells treated with puromycin (+) or untreated (-), the levels of β-actin and Coomassie staining as loading controls. (D) Translation rate determined by relative puromycin incorporation to β-actin. Statistical significance by Mann-Whitney U test, ** $p<0.01$. Values represent mean. Error bars represent SEM.

**Supplementary Figure 2: Interaction between nucleolar stress and mHTT accumulation.** (A) Quantification of the percentage of nuclei with nucleolar NPM1 and NCL localization analysed in the four experimental groups as mean values. (N, number of mice; control: 4, TIF-IA$^{D1RCre}$: 3, R6/2: 4, dm: 5). Statistical significance evaluated by Kruskal-Wallis test ($p=0.0005$) and post-hoc Dunn's multiple comparison; * $p<0.05$, ** $p<0.01$ (% nuclei with nucleolar NCL: controls vs. TIF-IA$^{D1Cre}$, $p=0.008$; controls vs. dm, $p=0.015$; TIF-IA$^{D1Cre}$ vs. R6/2, $p=0.017$; R6/2 vs. dm, $p=0.033$. Percentage of nuclei with nucleolar NPM1: controls vs. TIF-IA$^{D1Cre}$, $p=0.033$; controls vs. dm, $p=0.037$; TIF-IA$^{D1Cre}$ vs. R6/2, $p=0.22$; R6/2 vs. dm, $p=0.30$). (B) Quantification of the percentage of striatal nuclei with intranuclear mHTT inclusions and of their area in R6/2 and dm mice at 9 weeks. Diagrams show mean values error bars represent SEM (N, number of mice; R6/2: 4, dm: 4). (C, D) Representative images of immunohistochemistry performed on striatal sections by the EM48 antibody. (E) Nuclear distribution of mHTT measured as the percentage of nuclei with diffuse mHTT per field of view (FOV). Diagrams show mean values. Error bars represent SEM (N, number of mice; R6/2: 4, dm: 4). Statistical significance by Mann-Whitney U test ($p=0.03$). Arrows point to nuclei in insets as examples of nuclei with diffuse mHTT, while circled area shows an example of nuclei without mHTT in the nucleoplasm. Scale bar 100μm, inset 20μm.

**Supplementary Figure 3: Behavior analysis of pre-symptomatic zQ175 mice at 5 months and dysregulation of rRNA transcription and processing factors in the zQ175 mice.** (A) Body weight of zQ175 and control mice at 5 months shows no significant differences between control and mutant. (B) Endurance on an accelerating rotarod shows no significant differences at 5 months between control and zQ175 mice by Mann-Whitney U test (N: number of mice, control = 10, zQ175= 13). (C) Grip strength test at 5 months shows no differences between control and zQ175 mice. N: number of mice (in blue: males, in red: females), control = 10, zQ175= 13; values represent mean ±SEM. (D,E) The log2 fold changes of rRNA transcription and rRNA processing factors are shown for the analysis of human BA4 cortex RNAseq data (GSE79666)(Lin L *et al.*, 2016) and data from the striata and gastrocnemius of zQ175 mice at different ages (HDinHD.org)(Langfelder P *et al.*, 2016). rRNA transcription factors were chosen according to (Drygin D *et al.*, 2010; Sharifi S and Bierhoff H, 2018) and rRNA processing factors according to (Aubert M *et al.*, 2018). Only genes that were significantly changed (Benjamini-Hochberg adjusted *p*-value < 0.05) in at least one of the analysis datasets are shown. For the full list see Excel file.

**Supplementary Figure 4: No differences in the nuclear area in the skeletal muscle of zQ175 mice and HD patients, NPM1 signal area is not affected in the skeletal muscle of mutant mice showing striatal neurodegeneration.** (A, B) Mean area of DAPI stained nuclei at 5 and 10 months in muscle (quadriceps) of control (N=5, 4) and zQ175 (N=4, 6) mice. (C) No significant differences between the three groups (N= 5 for each group) in the nuclear area assessed by DAPI signal area. Values represent mean. Error bars represent SEM. (D) Representative confocal images of quadriceps cryosections stained for NPM1 (green) in control and TIF-IA$^{D1Cre}$ mice at 3 months. Nuclei are labelled with DAPI (blue). The arrows point out to NPM1 signal. Scale bar: 20µm. (E) Quantification of the percentage of nuclei with nucleolar localisation of NPM1 in control (N=6) and TIF-IA$^{D1Cre}$ mice (N= 4); p=0.0095 by Mann-Whitney U test. Mean area of the NPM1 signal (in µm$^2$) in control (N=5) and TIF-IA$^{D1Cre}$ (N= 4); p= 0.0317 by Mann-Whitney U test. Error bars represent SEM. *

p<0.05, ** p<0.01. (F) Correlation between manual and automated counting of NPM1 signals and area in human muscle biopsies from healthy controls and pre-/early Huntington's disease. Representative original and segmented images are shown for comparison. Scale bar: 10µm. Quantification of the percentage of nuclei showing nucleolar NPM1 signal by a machine learning algorithm (semantic convoluted neuronal network) in pre- and early-Huntington's disease individuals in comparison with age-matched controls (N=5 for each group). A significant decrease of NPM1 signal was confirmed in early Huntington's disease by Kruskal-Wallis test and Dunn's multiple comparison (p=0.002 early-Huntington's disease vs. controls). Pearson correlation between the manual and automated quantification is significant (p<0.0001). Mean area of the NPM1 signal (in $\mu m^2$) in control, pre- and early-Huntington's disease individuals (N=5 for each group). Statistical significance is assessed by Kruskal-Wallis test and Dunn's multiple comparison (p=0.027 early Huntington's disease vs. control). Pearson correlation between the manual and automated quantification is significant (p=0.0003). * p<0.05, ** p<0.01.

**Supplementary Table 1: Summary of the human cohorts for which quadriceps biopsies were analyzed.**

| Patient cohort | control | pre-HD | early HD |
|---|---|---|---|
| Age (years) | 38 ± 3.9 | 43.2 ± 9.1 | 43.4 ± 5.2 |
| Gender | 1 m / 4 f | 3 m /2 f | 2 m /3 f |
| Number of CAG repeats | n/a | 43.6 ± 1.5 | 45.2 ± 2.8 |
| DBS (=age*(CAG-35.5)) | n/a | 346 ± 73.4 | 420.5 ± 132.6 |

*Abbreviations in Table: m, male; f, female; DBS, disease burden score;* ± SD

**Supplementary Table 2: Overview of the functional and structural changes of the nucleolus by the analysis of in pre-rRNA synthesis and NPM1 and NCL immunostaining in different models of Huntington's disease and in human muscle biopsies.**

| | Q111/111 cells | R6/2 striatum pre-/sym | zQ175 striatum pre-sym | zQ175 quadriceps sym | HD quadriceps |
|---|---|---|---|---|---|
| **Pre-rRNA** | = [a)] | - [a) b)] | = | - | - [c)] |
| **NPM1 in nucleoli** | - | - [a)] | - | - | - |
| **NCL in nucleoli** | = | = | = | = | = |

***Abbreviations in Table:*** *pre-sym, pre-symptomatic; sym, symptomatic; HD, Huntington's disease;*
*-, decreased; +, increased; [a)] Lee J et al, 2011; [b)] Kreiner et al. 2013; [c)] Jesse et al. 2017.*

**Supplementary Table 3: List of mouse TaqMan assays**

| Gene | Assay ID | Primer/probe or consensus sequence | Gene –bank No. | Length (bp) | EB |
|---|---|---|---|---|---|
| *Rn18s* | Mm03928990_g1 | TACTTGGATAACTGTGGTAATTCTA | NR_003278.3 | 61 | - |
| *D1r* | Mm01353211_m1 | CCCAGATCGGGCATTTGGAGAGATG | NM_010076.3 | 65 | 1-2 |
| *D2r* | Mm00438541_m1 | GTCGTCTATCTGGAGGTGGTGGGTG | NM_010077.2 | 71 | 2-3 |
| *Hprt* | Mm01545399_m1 | GGACTGATTATGGACAGGACTGAAA | NM_013556.2 | 81 | 2-3 |
| *Metap1* | Mm00558361_m1 | ACTTCTGCTCGCAGGAATGCTTTAA | NM_175224.4 | 56 | 1-2 |

*Abbreviations: Length, amplicon length; EB: exon boundary*
