## Supplementary material for "Nucleolar stress controls mutant Huntingtin toxicity and monitors Huntington’s disease progression": Suppl stat information tables

### Supplementary statistical information

#### Table 1

Data and statistics for graphs shown in **Figure 1B**, n: number of nuclei, N: number of microscopic fields in two independent experiments. Statistical significance for NCL was assessed according to two-tailed t-test. Mann-Whitney U test (MWU) was not applicable due to lack of variance in the Q7/7 samples.

| **B) % nuclei with nucleolar NPM1** | | | | | | |
|---|---|---|---|---|---|---|
| | **mean** | **±SD** | **±SEM** | **median** | **95% CI** | **n/N** |
| **Q7/7** | 82.75 | 12.87 | 5.255 | 82.35 | 69.24- 96.25 | 421/6 |
| **Q111/111** | 59.26 | 14.38 | 5.871 | 56.74 | 44.17-74.35 | 634/6 |
| **MWU** | **0.0152** | | | | | |
| t-test | 0.0138 | | | | | |

| **B) % nuclei with nucleolar NCL** | | | | | | |
|---|---|---|---|---|---|---|
| | **Mean** | **±SD** | **±SEM** | **median** | **95% CI** | **n/N** |
| **Q7/7** | 100 | 0 | 0 | 100 | 100-100 | 180/11 |
| **Q111/111** | 95.45 | 7.778 | 3.175 | 98.21 | 87.29-103.6 | 135/6 |
| **t-test** | 0.0646 | | | | | |

#### Table 2

Data and statistics for graphs shown in **Suppl. Figure 1B, D**; N: total number of repeats performed respectively in four and two independent experiments. * Statistical significance is shown according to the Mann-Whitney U test (MWU).

| **B) polysomes / total ribosomes (% of control)** | | | | | | |
|---|---|---|---|---|---|---|
| | **Mean** | **±SD** | **±SEM** | **median** | **95% CI** | **N** |
| **Q7/7** | 100 | 9.03 | 3.19 | 100 | 92.5-107.6 | 8 |
| **Q111/111** | 81.99 | 15.3 | 5.41 | 78.6 | 69.2-94.8 | 8 |
| **MWU** | **0.038** | | | | | |
| **t-test** | 0.012 | | | | | |

| **D) puromycin incorporation (% of control)** | | | | | | |
|---|---|---|---|---|---|---|
| | **Mean** | **±SD** | **±SEM** | **median** | **95% CI** | **N** |
| **Q7/7** | 100 | 16.9 | 6.9 | 100.8 | 82.3-117.7 | 6 |
| **Q111/111** | 69.4 | 7.5 | 3.4 | 67.0 | 60.1-78.7 | 5 |
| **MWU** | **0.0087** | | | | | |
| **t-test** | 0.0047 | | | | | |

**Table 3**

Data and statistics for graphs shown in **Fig. 2B**, n: number of analysed striatal nuclei, N: number of mice. Statistical significance is shown according to the Mann-Whitney U test (MWU).

| **B) mHTT intensity (nucleoplasm / inclusion ratio)** | | | | | | |
|---|---|---|---|---|---|---|
| | **mean** | **±SD** | **±SEM** | **Median** | **95% CI** | **n/N** |
| **R6/2** | 0.25 | 0.03 | 0.01 | 0.25 | 0.20-0.29 | 335/4 |
| **dm** | 0.18 | 0.01 | 0.01 | 0.18 | 0.16-0.20 | 296/4 |
| ***MWU** | **0.0286** | | | | | |
| **t-test** | 0.0046 | | | | | |

**Table 4**

Data and statistics for graphs shown in **Suppl**. **Fig. 2**, n: number of analysed striatal nuclei, N: number of mice. Statistical significance is shown according to Kruskal-Wallis test (non-parametric one way analysis of variance).

| **A) % nuclei with nucleolar NCL** | | | | | | |
|---|---|---|---|---|---|---|
| | **Mean** | **±SD** | **±SEM** | **Median** | **95% CI** | **n/N** |
| **control** | 91.09 | 5.01 | 2.50 | 89.31 | 83.12-99.06 | 217/4 |
| **TIF-IA$^{D1Cre}$** | 10.23 | 4.28 | 2.47 | 9.524 | -0.39-20.85 | 185/3 |
| **R6/2** | 89.05 | 4.91 | 2.45 | 88.08 | 81.25-96.86 | 335/4 |
| **dm** | 15.36 | 5.67 | 2.53 | 13.68 | 8.322-22.4 | 371/5 |

| **Kruskal-Wallis test p=0.0005** |
|---|
| **Dunn's multiple comparison** |
| **control vs. TIF-IA$^{D1Cre}$ 0.0079, control vs. R6/2** 0.7664, **control vs. dm 0.0146, TIF-IA$^{D1Cre}$ vs. R6/2 0.0172, TIF-IA$^{D1Cre}$ vs. dm** 0.5914, **R6/2 vs. dm 0.0332** |

| **A) % nuclei with nucleolar NPM1** | | | | | | |
|---|---|---|---|---|---|---|
| | **Mean** | **±SD** | **±SEM** | **Median** | **95% CI** | **n/N** |
| **control** | 73.2 | 11.8 | 5.9 | 69.5 | 54.5-92 | 280/4 |
| **TIF-IA$^{D1Cre}$** | 2.9 | 0.7 | 0.4 | 3.2 | 1.1-4.7 | 241/3 |
| **R6/2** | 59.5 | 15.5 | 7.8 | 55.4 | 34.8-84.2 | 343/4 |
| **dm** | 3.6 | 1.6 | 0.7 | 3.9 | 1.5-5.6 | 464/5 |

| **Kruskal-Wallis test p=0.0003** |
|---|
| **Dunn's multiple comparison** |
| **control vs. TIF-IA$^{D1Cre}$ 0.0333, control vs. R6/2** >0.9999, **control vs. dm 0.0369, TIF-IA$^{D1Cre}$ vs. R6/2** 0.2221, **TIF-IA$^{D1Cre}$ vs. dm** >0.9999, **R6/2 vs. dm** 0.3021 |

| **B) Intranuclear mHTT inclusions (% nuclei)** | | | | | | |
|---|---|---|---|---|---|---|
| | **Mean** | **±SD** | **±SEM** | **Median** | **95% CI** | **n/N** |
| **R6/2** | 85.20 | 9.44 | 4.72 | 87.03 | 70.18-100.2 | 335/4 |
| **dm** | 88.97 | 3.16 | 1.58 | 89.07 | 83.94-93.99 | 296/4 |
| **MWU** | 0.8857 | | | | | |
| **t-test** | 0.4781 | | | | | |

| B) mHTT inclusion area (µm²) | | | | | | |
|---|---|---|---|---|---|---|
| | **Mean** | **±SD** | **±SEM** | **Median** | **95% CI** | **n/N** |
| **R6/2** | 3.18 | 0.44 | 0.22 | 3.296 | 2.49- 3.88 | 335/4 |
| **dm** | 3.66 | 0.18 | 0.09 | 3.614 | 3.38-3.94 | 296/4 |
| **MWU** | 0.1143 | | | | | |
| **t-test** | 0.0894 | | | | | |

| E) Nuclei with diffuse mHTT/field of view (FOV) (%) | | | | | | |
|---|---|---|---|---|---|---|
| | **mean** | **±SD** | **±SEM** | **Median** | **95% CI** | **n/N** |
| **R6/2** | 82.38 | 15.56 | 7.78 | 86.00 | 57.6- 107.1 | 575/4 |
| **dm** | 17.50 | 4.80 | 2.40 | 16.50 | 9.87-25.1 | 584/4 |
| **MWU** | **0.0286** | | | | | |
| **t-test** | 0.0002 | | | | | |

**Table 5**

Data and statistics for body weight of mice for behaviour analysis and for the graphs shown in **Figure 2D, E**, N: number of mice, w: weeks. * Statistical significance is shown according to Kruskal-Wallis test (non-parametric one way analysis of variance) or two-way ANOVA.

| Body weight (9w) (% of control) | | | | | | |
|---|---|---|---|---|---|---|
| | **mean** | **±SD** | **±SEM** | **Median** | **95% CI** | **N (m/f)** |
| **Control** | 100 | 5.7 | 2.0 | 100.6 | 95.2-104.8 | 8 (4/4) |
| **TIF-IA$^{D1Cre}$** | 96.3 | 4.9 | 2.0 | 96.4 | 91.2-101.4 | 6 (2/4) |
| **R6/2** | 99.7 | 8.7 | 2.5 | 99.3 | 94.2-105.2 | 12 (5/7) |
| **dm** | 95.8 | 6.6 | 1.9 | 95.7 | 91.6-99.6 | 12 (7/5) |
| ***Kruskal-Wallis test** p=0.424 | | | | | | |
| **p values from Dunn's multiple comparisons**<br>**Control vs. TIF-IA$^{D1Cre}$** >0.9999, **Control vs. R6/2** >0.9999, **Control vs. dm** =0.734, **TIF-IA$^{D1Cre}$ vs. R6/2** >0.9999, **TIF-IA$^{D1Cre}$ vs. dm** >0.9999, **R6/2 vs. dm** >0.9999 | | | | | | |

| D) Rotarod endurance (s) | | | | |
|---|---|---|---|---|
| | | **trial 1**<br>**9 w** | **trial 2**<br>**10 w** | **trial 3**<br>**11 w** |
| **Control** | **Mean** | 358.8 | 366.2 | 391.8 |
| | **±SD** | 57.2 | 66.1 | 73.0 |
| | **±SEM** | 20.2 | 23.4 | 25.8 |
| | **Median** | 353.3 | 364.3 | 392.5 |
| | **upper limit** | 463.0 | 480.0 | 480.0 |
| | **lower limit** | 287.0 | 284.3 | 285.0 |
| | **N (m/f)** | 8 (4/4) | 8 (4/4) | 8 (4/4) |
| **TIF-IA$^{D1Cre}$** | **Mean** | 372.5 | 356.8 | 416.7 |
| | **±SD** | 90.1 | 119.4 | 58.8 |
| | **±SEM** | 36.8 | 48.8 | 24.0 |
| | **Median** | 380.2 | 347.0 | 424.0 |
| | **upper limit** | 470.0 | 513.3 | 480.0 |
| | **lower limit** | 242.0 | 167.3 | 313.0 |
| | **N (m/f)** | 6 (2/4) | 6 (2/4) | 6 (2/4) |
| **R6/2** | **Mean** | 252.8 | 195.9 | 191.1 |
| | **±SD** | 87.0 | 69.0 | 70.8 |

| | | | | |
|---|---|---|---|---|
| | **±SEM** | 25.1 | 19.9 | 20.4 |
| | **Median** | 242.7 | 202.2 | 188.3 |
| | **upper limit** | 413.7 | 296.0 | 283.3 |
| | **lower limit** | 140.0 | 72.0 | 52.0 |
| | **N (m/f)** | 12 (5/7) | 12 (5/7) | 12 (5/7) |
| **dm** | **Mean** | 196.1 | 108.2 | 92.1 |
| | **±SD** | 61.5 | 33.6 | 33.4 |
| | **±SEM** | 17.8 | 9.7 | 9.6 |
| | **Median** | 200.7 | 115.5 | 96.0 |
| | **upper limit** | 293.0 | 162.5 | 141.0 |
| | **lower limit** | 92.7 | 46.3 | 36.5 |
| | **N (m/f)** | 12 (7/5) | 12 (7/5) | 12 (7/5) |

| ***Two-way ANOVA** | |
|---|---|
| **Source of Variation** | **P value** |
| **Interaction** | **0.0222** |
| **Age** | **0.0681** |
| **Genotype** | **<0.0001** |

| **Results of Tukey's multiple comparison p value** | **trial 1** | **trial 2** | **trial 3** |
|---|---|---|---|
| **Control vs. TIF-IA$^{D1Cre}$** | 0.9824 | 0.9942 | 0.9069 |
| **Control vs. R6/2** | **0.0053** | **<0.0001** | **<0.0001** |
| **Control vs. dm** | **<0.0001** | **<0.0001** | **<0.0001** |
| **TIF-IA$^{D1Cre}$ vs. R6/2** | **0.0038** | **<0.0001** | **<0.0001** |
| **TIF-IA$^{D1Cre}$ vs. dm** | **<0.0001** | **<0.0001** | **<0.0001** |
| **R6/2 vs. dm** | 0.1836 | **0.0117** | **0.0033** |

| E) Grip strength (mN) | | | | |
|---|---|---|---|---|
| | | trial 1<br>9 w | trial 2<br>10 w | trial 3<br>11 w |
| Control | Mean strength (mN) | 616.7 | 641.4 | 652.7 |
| | ±SD | 89.6 | 101.3 | 95.4 |
| | ±SEM | 31.7 | 35.8 | 33.7 |
| | Median | 599.2 | 670.2 | 670.5 |
| | upper limit | 793.7 | 742.7 | 782.7 |
| | lower limit | 501.7 | 449.7 | 514.3 |
| | N (m/f) | 8 (4/4) | 8 (4/4) | 8 (4/4) |
| TIF-IA$^{D1Cre}$ | Mean strength (mN) | 693.5 | 627.7 | 660.3 |
| | ±SD | 156.3 | 124.5 | 81.1 |
| | ±SEM | 63.8 | 50.8 | 33.1 |
| | Median | 720.7 | 678.3 | 644.0 |
| | upper limit | 858.0 | 708.3 | 769.7 |
| | lower limit | 406.7 | 380.0 | 564.7 |
| | N (m/f) | 6 (2/4) | 6 (2/4) | 6 (2/4) |
| R6/2 | Mean strength (mN) | 700.6 | 640.2 | 580.6 |
| | ±SD | 119.2 | 120.2 | 94.1 |
| | ±SEM | 34.4 | 34.7 | 27.2 |
| | Median | 738.8 | 629.7 | 611.5 |
| | upper limit | 838.0 | 817.7 | 688.3 |
| | lower limit | 413.3 | 494.0 | 408.3 |
| | N (m/f) | 12 (5/7) | 12 (5/7) | 12 (5/7) |
| dm | Mean strength (mN) | 616.4 | 415.2 | 333.3 |
| | ±SD | 103.2 | 95.5 | 104.0 |
| | ±SEM | 29.8 | 27.6 | 30.0 |
| | Median | 648.7 | 413.3 | 341.0 |
| | upper limit | 750.3 | 581.0 | 463.7 |
| | lower limit | 409.7 | 232.7 | 181.7 |
| | N (m/f) | 12 (7/5) | 12 (7/5) | 12 (7/5) |

| Two-way ANOVA | |
|---|---|
| Source of Variation | P value |
| Interaction | 0.0006 |
| Age | 0.0005 |
| Genotype | <0.0001 |

| Results of Tukey's multiple comparison p values | trial 1 | trial 2 | trial 3 |
|---|---|---|---|
| Control vs. TIF-IA$^{D1Cre}$ | 0.5484 | 0.9954 | 0.9992 |
| Control vs. R6/2 | 0.3215 | >0,9999 | 0.4581 |
| Control vs. Dm | >0,9999 | **<0,0001** | **<0,0001** |
| TIF-IA$^{D1Cre}$ vs. R6/2 | 0.9992 | 0.9955 | 0.4497 |
| TIF-IA$^{D1Cre}$ vs. dm | 0.4786 | **0.0008** | **<0,0001** |
| R6/2 vs. dm | 0.2245 | **<0,0001** | **<0,0001** |

**Table 6**

Data and statistics for graphs shown in **Fig. 3,** N: number of mice. * Statistical significance is shown according to the Mann-Whitney U test (MWU).

| **A) *D2r* qPCR** | | | | | | |
|---|---|---|---|---|---|---|
| | | **3 mo** | **4 mo** | **5 mo** | **6 mo** | **10 mo** |
| **control** | **mean** | 1 | 1 | 1 | 1 | 1 |
| | **±SD** | 0.34 | 0.34 | 0.23 | 0.38 | 0.29 |
| | **±SEM** | 0.14 | 0.11 | 0.09 | 0.12 | 0.13 |
| | **median** | 1.00 | 0.98 | 0.91 | 0.96 | 0.96 |
| | **upper limit** | 1.40 | 1.69 | 1.26 | 1.50 | 1.34 |
| | **lower limit** | 0.67 | 0.66 | 0.72 | 0.41 | 0.67 |
| | **N** | 6 | 9 | 7 | 10 | 5 |
| **zQ175** | **mean** | 1.25 | 0.72 | 0.63 | 0.51 | 0.58 |
| | **±SD** | 0.21 | 0.12 | 0.20 | 0.19 | 0.25 |
| | **±SEM** | 0.09 | 0.04 | 0.08 | 0.07 | 0.11 |
| | **median** | 1.28 | 0.77 | 0.59 | 0.48 | 0.61 |
| | **upper limit** | 1.45 | 0.81 | 0.99 | 0.84 | 0.91 |
| | **lower limit** | 0.93 | 0.46 | 0.43 | 0.22 | 0.26 |
| | **N** | 5 | 8 | 6 | 8 | 5 |
| ***MWU** | | 0.178 | 0.114 | **0.014** | **0.012** | **0.032** |
| **t-test** | | 0.186 | **0.037** | **0.010** | **0.005** | **0.036** |

| **A) *D1r* qPCR** | | | | | | |
|---|---|---|---|---|---|---|
| | | **3 mo** | **4 mo** | **5 mo** | **6 mo** | **10 mo** |
| **control** | **mean** | 1 | 1 | 1 | 1 | 1 |
| | **±SD** | 0.19 | 0.39 | 0.17 | 0.47 | 0.19 |
| | **±SEM** | 0.08 | 0.13 | 0.07 | 0.15 | 0.09 |
| | **median** | 0.96 | 0.86 | 1.02 | 0.97 | 0.95 |
| | **upper limit** | 1.28 | 1.90 | 1.21 | 1.76 | 1.31 |
| | **lower limit** | 0.78 | 0.60 | 0.70 | 0.34 | 0.79 |
| | **N** | 6 | 9 | 7 | 10 | 5 |
| **zQ175** | **mean** | 0.96 | 0.97 | 0.89 | 0.66 | 0.75 |
| | **±SD** | 0.34 | 0.10 | 0.18 | 0.10 | 0.11 |
| | **±SEM** | 0.14 | 0.04 | 0.07 | 0.04 | 0.05 |
| | **median** | 0.88 | 0.96 | 0.90 | 0.66 | 0.68 |
| | **upper limit** | 1.55 | 1.10 | 1.18 | 0.85 | 0.89 |
| | **lower limit** | 0.56 | 0.83 | 0.66 | 0.52 | 0.64 |
| | **N** | 6 | 8 | 6 | 8 | 5 |
| ***MWU** | | 0.699 | 0.541 | 0.295 | 0.101 | **0.032** |
| **t-test** | | 0.828 | 0.825 | 0.270 | 0.061 | **0.038** |

| **D) intranuclear mHTT inclusions (% nuclei)** | | | | | | |
|---|---|---|---|---|---|---|
| | **mean** | **±SD** | **±SEM** | **Median** | **95% CI** | **n/N** |
| **5 months** | 33.29 | 17.36 | 7.765 | 36.11 | 11.74-54.85 | 409/5 |
| **10 months** | 76.89 | 6.242 | 2.791 | 78.61 | 69.14-84.64 | 416/5 |
| **MWU** | | **0.0079** | | | | |
| **t-test** | | **0.0007** | | | | |

| E) mHTT inclusion area (µm²) | | | | | | |
|---|---|---|---|---|---|---|
| | **mean** | **±SD** | **±SEM** | **Median** | **95% CI** | **n/N** |
| **5 months** | 1.21 | 0.21 | 0.09 | 1.17 | 0.96-1.47 | 409/5 |
| **10 months** | 1.61 | 0.33 | 0.15 | 1.74 | 1.21-2.02 | 416/5 |
| **MWU** | | 0.0952 | | | | |
| **t-test** | | **0.0487** | | | | |

| F) mHTT intensity nucleoplasm/ inclusion ratio | | | | | | |
|---|---|---|---|---|---|---|
| | **mean** | **±SD** | **±SEM** | **Median** | **95% CI** | **n/N** |
| **5 months** | 0.64 | 0.11 | 0.05 | 0.61 | 0.51-0.78 | 409/5 |
| **10 months** | 0.44 | 0.05 | 0.02 | 0.43 | 0.38-0.50 | 416/5 |
| **MWU** | | **0.0159** | | | | |
| **t-test** | | **0.0054** | | | | |

**Table 7**

Data and statistics for graphs shown in **Suppl. Figure 3**; N: number of mice. Statistical significance is shown according to the Mann-Whitney U test (MWU).

| A) Body weight (% of control) | | | | | | | |
|---|---|---|---|---|---|---|---|
| | **Mean** | **±SD** | **±SEM** | **Median** | **95% CI** | **N** | **m/f** |
| **Control** | 100 | 8.2 | 2.5 | 103.2 | 94.5-105.5 | 11 | 5/6 |
| **zQ175** | 96.9 | 7.6 | 2.1 | 96.9 | 92.3-101.5 | 13 | 6/7 |
| **MWU** | 0.3607 | | | | | | |
| **t-test** | 0.3464 | | | | | | |

| B) Rotarod endurance (s) | | | | | | | |
|---|---|---|---|---|---|---|---|
| | **Mean** | **±SD** | **±SEM** | **Median** | **95% CI** | **N** | **m/f** |
| **Control** | 381 | 83.97 | 26.55 | 406.3 | 320.9-441 | 10 | 4/6 |
| **zQ175** | 324.7 | 83.86 | 23.26 | 282 | 274-375.4 | 13 | 6/7 |
| **MWU** | 0.1346 | | | | | | |
| **t-test** | 0.1259 | | | | | | |

| C) Grip strength (mN) | | | | | | | |
|---|---|---|---|---|---|---|---|
| | **mean** | **±SD** | **±SEM** | **Median** | **95% CI** | **N** | **m/f** |
| **control** | 541.8 | 149.6 | 45.12 | 586.3 | 441.2-642.3 | 11 | 5/6 |
| **zQ175** | 525.1 | 104.2 | 28.9 | 524.7 | 462.1-588.1 | 13 | 6/7 |
| **MWU** | 0.4244 | | | | | | |
| **t-test** | 0.7513 | | | | | | |

**Table 8**

Data and statistics for graphs shown in **Figure 4**, n: number of analysed striatal cells, N: number of mice. * Statistical significance is shown according to the Mann-Whitney U test (MWU).

| B) 47S1 pre-rRNA | | | |
|---|---|---|---|
| | | **5 months** | **10 months** |
| **control** | **Mean (fold change relative to control)** | 1 | 1 |
| | **±SD** | 0.19 | 0.22 |
| | **±SEM** | 0.07 | 0.09 |
| | **Median** | 0.96 | 0.95 |
| | **upper limit** | 1.28 | 1.41 |
| | **lower limit** | 0.77 | 0.82 |
| | **N** | 7 | 6 |
| **zQ175** | **Mean (fold change relative to control)** | 0.89 | 1.15 |
| | **±SD** | 0.25 | 0.11 |
| | **±SEM** | 0.10 | 0.06 |
| | **Median** | 0.84 | 1.14 |
| | **Upper 95% CI** | 1.23 | 1.29 |
| | **Lower 95% CI** | 0.51 | 1.03 |
| | **N** | 6 | 4 |
| **MWU** | | 0.366 | 0.2571 |
| **t-test** | | 0.3836 | 0.2476 |

| C) 47S2 pre-rRNA | | | |
|---|---|---|---|
| | | **5 months** | **10 months** |
| **Control** | **Mean (fold change relative to control)** | 1 | 1 |
| | **±SD** | 0.39 | 0.42 |
| | **±SEM** | 0.15 | 0.17 |
| | **Median** | 1.11 | 0.84 |
| | **upper limit** | 1.37 | 1.77 |
| | **lower limit** | 0.29 | 0.65 |
| | **N** | 7 | 6 |
| **zQ175** | **Mean (fold change relative to control)** | 0.69 | 1.20 |
| | **±SD** | 0.28 | 0.56 |
| | **±SEM** | 0.13 | 0.25 |
| | **median** | 0.56 | 1.20 |
| | **Upper 95% CI** | 1.10 | 2.00 |
| | **Lower 95% CI** | 0.43 | 0.49 |
| | **N** | 5 | 5 |
| **MWU** | | 0.8238 | 0.4286 |
| **t-test** | | 0.8092 | 0.5228 |

| D) 18S rRNA | | | |
|---|---|---|---|
| | | **5 months** | **10 months** |
| **Control** | **Mean (fold change relative to control)** | 1 | 1 |
| | **±SD** | 0.23 | 0.06 |
| | **±SEM** | 0.08 | 0.03 |
| | **Median** | 0.96 | 0.99 |
| | **upper limit** | 1.39 | 1.08 |
| | **lower limit** | 0.73 | 0.91 |
| | **N** | 8 | 6 |

| | | | |
|---|---|---|---|
| **zQ175** | **Mean (fold change relative to control)** | 1.00 | 1.07 |
| | **±SD** | 0.20 | 0.29 |
| | **±SEM** | 0.08 | 0.13 |
| | **median** | 0.98 | 0.95 |
| | **Upper 95% CI** | 1.29 | 1.39 |
| | **Lower 95% CI** | 0.74 | 0.81 |
| | **N** | 6 | 5 |
| **MWU** | | 0.9497 | 0.7922 |
| **t-test** | | 0.9748 | 0.558 |

| **F) pre-rRNA foci per cell** | | | |
|---|---|---|---|
| **Age** | | **5 months** | **10 months** |
| **Control** | **Mean (fold change relative to control)** | 1 | 1 |
| | **±SD** | 0.12 | 0.21 |
| | **±SEM** | 0.04 | 0.08 |
| | **median** | 1.00 | 1.01 |
| | **upper limit** | 0.91 | 0.80 |
| | **lower limit** | 1.09 | 1.20 |
| | **n** | 7793 | 4841 |
| | **N** | 9 | 7 |
| **zQ175** | **Mean (fold change relative to control)** | 1.19 | 0.97 |
| | **±SD** | 0.23 | 0.15 |
| | **±SEM** | 0.09 | 0.06 |
| | **median** | 1.09 | 0.99 |
| | **Upper 95% CI** | 0.98 | 0.81 |
| | **Lower 95% CI** | 1.40 | 1.12 |
| | **n** | 6494 | 5001 |
| | **N** | 7 | 6 |
| **MWU** | | 0.0907 | 0.6282 |
| **t-test** | | **0.0441** | 0.771 |

| **G) 5.8S / 5S rRNA by Northern blot** | | | | | | |
|---|---|---|---|---|---|---|
| | **mean** | **±SD** | **±SEM** | **median** | **95% CI** | **N** |
| **Control** | 1.119 | 0.3084 | 0.09751 | 1.036 | 0.898-1.34 | 10 |
| **zQ175** | 1.155 | 0.2881 | 0.1018 | 1.051 | 0.914-1.396 | 8 |
| **MWU** | 0.6965 | | | | | |
| **t-test** | 0.8057 | | | | | |

**Table 9**

Data and statistics for graphs shown in **Fig. 5C**, n: number of analysed striatal nuclei, N: number of mice. * Statistical significance is shown according to the Mann-Whitney U test (MWU).

| **5 months: % of nuclei with nucleolar NPM1** | | | | | | | | |
|---|---|---|---|---|---|---|---|---|
| | **mean** | **±SD** | **±SEM** | **median** | **95% CI** | **N/n** | ***MWU** | **t-test** |
| **control** | 66.3 | 10.3 | 3.9 | 67.0 | 56.7-75.8 | 7/724 | **0.0205** | **0.0108** |
| **zQ175** | 41.6 | 19.7 | 7.0 | 42.2 | 25.1-58.0 | 8/720 | | |

| **5 months: % of nuclei with nucleolar NCL** | | | | | | | | |
|---|---|---|---|---|---|---|---|---|
| | **mean** | **±SD** | **±SEM** | **median** | **95% CI** | **N/n** | **MWU** | **t-test** |
| **control** | 84.1 | 8.2 | 3.1 | 82.7 | 76.6-91.7 | 7/480 | 0.7551 | 0.7489 |
| **zQ175** | 85.5 | 4.6 | 2.1 | 85.5 | 79.7-91.3 | 5/409 | | |

| **10 months: % of nuclei with nucleolar NPM1** | | | | | | | | |
|---|---|---|---|---|---|---|---|---|
| | **mean** | **±SD** | **±SEM** | **median** | **95% CI** | **N/n** | **MWU** | **t-test** |
| **control** | 77.7 | 14.9 | 6.7 | 70.7 | 59.2-96.3 | 5/771 | 0.1508 | 0.0756 |
| **zQ175** | 60.6 | 11.4 | 5.1 | 61.6 | 46.4-74.8 | 5/749 | | |

| **10 months: % of nuclei with nucleolar NCL** | | | | | | | | |
|---|---|---|---|---|---|---|---|---|
| | **mean** | **±SD** | **±SEM** | **median** | **95% CI** | **N/n** | **MWU** | **t-test** |
| **control** | 80.5 | 11.0 | 6.3 | 76.7 | 53.2-107.8 | 3/416 | >0.9999 | 0.9876 |
| **zQ175** | 80.4 | 8.0 | 3.6 | 80.7 | 70.5-90.2 | 5/387 | | |

**Table 10**

Data and statistics for graphs shown in **Fig. 6B-D**, n: number of DAPI positive nuclei in mouse quadriceps, N: number of mice. * Statistical significance is shown according to the Mann-Whitney U test (MWU).

| **B) 5 months: % nuclei with nucleolar NPM1** | | | | | | | | |
|---|---|---|---|---|---|---|---|---|
| | **mean** | **±SD** | **±SEM** | **median** | **95% CI** | **N/n** | **MWU** | **t-test** |
| **Control** | 33.91 | 4.90 | 2.45 | 35.47 | 26.13-41.70 | 4/269 | 0.4127 | 0.4862 |
| **zQ175** | 29.67 | 10.56 | 4.72 | 33.17 | 16.57-42.78 | 5/593 | | |

| **B) 5 months: % nuclei with nucleolar NCL** | | | | | | | | |
|---|---|---|---|---|---|---|---|---|
| | **mean** | **±SD** | **±SEM** | **Median** | **95% CI** | **N/n** | **MWU** | **t-test** |
| **Control** | 55.90 | 15.74 | 9.09 | 53.29 | 16.79-95.00 | 3/234 | >0.9999 | 0.7856 |
| **zQ175** | 59.33 | 13.04 | 7.53 | 63.04 | 26.94-91.72 | 3/250 | | |

| **B) 10 months: % nuclei with nucleolar NPM1** | | | | | | | | |
|---|---|---|---|---|---|---|---|---|
| | **mean** | **±SD** | **±SEM** | **median** | **95% CI** | **N/n** | **MWU** | **t-test** |
| **Control** | 44.57 | 12.58 | 6.291 | 48.74 | 24.55-64.59 | 4/516 | 0.2857 | 0.2698 |
| **zQ175** | 35.00 | 11.38 | 5.091 | 31.89 | 20.86-49.13 | 5/659 | | |

| **B) 10 months % nuclei with nucleolar NCL** | | | | | | | | |
|---|---|---|---|---|---|---|---|---|
| | **mean** | **±SD** | **±SEM** | **median** | **95% CI** | **N/n** | **MWU** | **t-test** |
| **control** | 53.54 | 7.16 | 3.58 | 54.38 | 42.15-64.94 | 4/750 | 0.7619 | 0.8672 |
| **zQ175** | 54.45 | 8.68 | 3.54 | 52.60 | 45.34-63.56 | 6/949 | | |

| **C) 5 months: NPM1 signal area (μm$^2$)** | | | | | | | | |
|---|---|---|---|---|---|---|---|---|
| | **mean** | **±SD** | **±SEM** | **Median** | **95% CI** | **N/n** | **MWU** | **t-test** |
| **control** | 3.16 | 0.51 | 0.228 | 2.927 | 2.53-3.79 | 5/220 | 0.2857 | 0.3456 |
| **zQ175** | 3.52 | 0.56 | 0.280 | 3.417 | 2.63-4.41 | 4/218 | | |

| **C) 5 months: NCL signal area (μm$^2$)** | | | | | | | | |
|---|---|---|---|---|---|---|---|---|
| | **Mean** | **±SD** | **±SEM** | **Median** | **95% CI** | **N/n** | **MWU** | **t-test** |
| **control** | 6.76 | 1.18 | 0.68 | 6.43 | 3.83-9.70 | 3/234 | 0.7000 | 0.4397 |
| **zQ175** | 5.84 | 1.45 | 0.84 | 6.26 | 2.25-9.43 | 3/250 | | |

| **C) 10 months NPM1 area (μm$^2$)** | | | | | | | | |
|---|---|---|---|---|---|---|---|---|
| | **mean** | **±SD** | **±SEM** | **median** | **95% CI** | **N/n** | ***MWU** | **t-test** |
| **Control** | 4.01 | 0.89 | 0.45 | 3.91 | 2.59-5.43 | 4/269 | **0.0381** | **0.0134** |
| **zQ175** | 2.67 | 0.47 | 0.19 | 2.69 | 2.18-3.16 | 6/306 | | |

| **C) 10 months NCL area (μm$^2$)** | | | | | | | | |
|---|---|---|---|---|---|---|---|---|
| | **mean** | **±SD** | **±SEM** | **median** | **95% CI** | **N/n** | **MWU** | **t-test** |
| **Control** | 5.66 | 1.17 | 0.59 | 5.24 | 3.80-7.52 | 4/269 | 0.4127 | 0.7006 |
| **zQ175** | 5.90 | 0.58 | 0.26 | 6.02 | 5.18-6.61 | 5/593 | | |

| D) 47S1 pre-rRNA muscle | | | |
|---|---|---|---|
| | | 5 months | 10 months |
| control | Mean (fold change relative to control) | 1 | 1 |
| | ±SD | 0.62 | 0.40 |
| | ±SEM | 0.36 | 0.20 |
| | Median | 0.83 | 0.95 |
| | upper limit | 1.69 | 1.48 |
| | lower limit | 0.48 | 0.61 |
| | N | 3 | 4 |
| zQ175 | Mean (fold change relative to control) | 0.83 | 0.79 |
| | ±SD | 0.43 | 0.36 |
| | ±SEM | 0.19 | 0.14 |
| | Median | 0.76 | 0.81 |
| | Upper 95% CI | 1.46 | 1.20 |
| | Lower 95% CI | 0.31 | 0.21 |
| | N | 5 | 7 |
| MWU | | 0.786 | 0.649 |
| t-test | | 0.667 | 0.401 |

| D) 47S2 pre-rRNA muscle | | | |
|---|---|---|---|
| | | 5 months | 10 months |
| Control | Mean (fold change relative to control) | 1 | 1 |
| | ±SD | 0.34 | 0.31 |
| | ±SEM | 0.20 | 0.15 |
| | Median | 1.05 | 0.98 |
| | upper limit | 1.31 | 1.39 |
| | lower limit | 0.64 | 0.65 |
| | N | 3 | 4 |
| zQ175 | Mean (fold change relative to control) | 0.96 | 0.54 |
| | ±SD | 0.45 | 0.13 |
| | ±SEM | 0.20 | 0.05 |
| | median | 0.85 | 0.53 |
| | Upper 95% CI | 1.58 | 0.70 |
| | Lower 95% CI | 0.38 | 0.32 |
| | N | 5 | 7 |
| *MWU | | >0,9999 | **0.012** |
| t-test | | 0.908 | **0.007** |

| D) 18S rRNA muscle | | | |
|---|---|---|---|
| | | 5 months | 10 months |
| Control | Mean (fold change relative to control) | 1 | 1 |
| | ±SD | 0.12 | 0.09 |
| | ±SEM | 0.06 | 0.05 |
| | Median | 0.98 | 1.02 |
| | upper limit | 1.19 | 1.15 |
| | lower limit | 0.81 | 0.85 |
| | N | 4 | 4 |
| zQ175 | Mean (fold change relative to control) | 1.16 | 1.03 |
| | ±SD | 0.26 | 0.23 |
| | ±SEM | 0.12 | 0.09 |
| | median | 1.25 | 1.08 |
| | Upper 95% CI | 1.48 | 1.24 |
| | Lower 95% CI | 0.84 | 0.82 |

| | | | |
|---|---|---|---|
| | **N** | 5 | 7 |
| **MWU** | | 0.286 | 0.788 |
| **t-test** | | 0.301 | 0.799 |

**Table 11**

Data and statistics for graphs shown in **Fig. 7B, C**, n: number of DAPI positive muscle nuclei, N: number of control individuals and Huntington's disease (HD) patients. * Statistical significance is shown according to the Kruskal-Wallis test.

| **B) % nuclei with nucleolar NPM1** | | | | | | |
|---|---|---|---|---|---|---|
| | **Mean** | **±SD** | **±SEM** | **median** | **95% CI** | **N/n** |
| **control** | 36.99 | 4.86 | 2.17 | 38.01 | 30.96-43.03 | 5/991 |
| **pre-HD** | 25.80 | 3.66 | 1.64 | 27.34 | 21.26-30.35 | 5/844 |
| **early HD** | 20.99 | 2.63 | 1.17 | 20.89 | 17.73-24.25 | 5/794 |
| **Kruskal-Wallis test p=0.0001** | | | | | | |
| **p values from Dunn's multiple comparison**<br>**control vs. pre-HD** 0.1431, **control vs. early HD 0.0027, pre-HD vs. early HD** 0.5373 | | | | | | |

| **B) % nuclei with nucleolar NCL** | | | | | | |
|---|---|---|---|---|---|---|
| | **Mean** | **±SD** | **±SEM** | **Median** | **95% CI** | **N/n** |
| **control** | 65.08 | 5.79 | 2.89 | 66.55 | 55.87-74.29 | 4/101 |
| **pre-HD** | 64.18 | 4.97 | 2.22 | 62.38 | 58.01-70.36 | 5/145 |
| **early HD** | 68.31 | 4.09 | 2.04 | 69.31 | 61.8-74.81 | 4/120 |
| **Kruskal-Wallis test** p=0.4499 | | | | | | |
| **p values from Dunn's multiple comparison**<br>**control vs. pre-HD>**0.9999, **control vs. early HD>**0.9999, **pre-HD vs. early HD** 0.5793 | | | | | | |

| **C) NPM1 area ($\mu m^2$)** | | | | | | |
|---|---|---|---|---|---|---|
| | **Mean** | **±SD** | **±SEM** | **median** | **95% CI** | **N/n** |
| **control** | 3.54 | 0.70 | 0.31 | 3.47 | 2.66-4.41 | 5/991 |
| **pre-HD** | 2.77 | 0.38 | 0.17 | 2.86 | 2.31-3.24 | 5/844 |
| **early HD** | 2.32 | 0.34 | 0.15 | 2.45 | 1.90-2.74 | 5/794 |
| **Kruskal-Wallis test p=0.0059** | | | | | | |
| **p values from Dunn's multiple comparison**<br>**control vs. pre-HD** 0.4708, **control vs. early HD 0.0099, pre-HD vs. early HD** 0.3843 | | | | | | |

| **C) NCL area ($\mu m^2$)** | | | | | | |
|---|---|---|---|---|---|---|
| | **Mean** | **±SD** | **±SEM** | **median** | **95% CI** | **N/n** |
| **control** | 2.73 | 0.60 | 0.30 | 2.69 | 1.77-3.69 | 4/101 |
| **pre-HD** | 2.78 | 0.63 | 0.28 | 2.44 | 2.00-3.55 | 5/145 |
| **early HD** | 2.69 | 0.41 | 0.21 | 2.57 | 2.03-3.35 | 4/120 |
| **Kruskal-Wallis test** p=0.8755 | | | | | | |
| **p values from Dunn's multiple comparison**<br>**control vs. pre-HD>**0.9999, **control vs. early HD>**0.9999, **pre-HD vs. early HD>**0.9999 | | | | | | |

**Table 12**

Data and statistics for graphs shown in **Suppl. Figure 4A-E**, n: number of DAPI positive nuclei in mouse quadriceps, N: number of mice. * Statistical significance is shown according to the Mann-Whitney U test (MWU) or Kruskal-Wallis test.

| **A) 5 months DAPI area (µm$^2$)** | | | | | | | | |
|---|---|---|---|---|---|---|---|---|
| | **mean** | **±SD** | **±SEM** | **Median** | **95% CI** | **N/n** | **MWU** | **t-test** |
| **control** | 27.51 | 4.714 | 2.108 | 27.15 | 21.66-33.36 | 5/607 | 0.4127 | 0.4093 |
| **zQ175** | 25.03 | 3.428 | 1.714 | 24.49 | 19.58-30.49 | 4/545 | | |

| **B) 10 months DAPI area (µm$^2$)** | | | | | | | | |
|---|---|---|---|---|---|---|---|---|
| | **mean** | **±SD** | **±SEM** | **Median** | **95% CI** | **N/n** | **MWU** | **t-test** |
| **control** | 21.92 | 2.26 | 1.13 | 21.66 | 18.32-25.53 | 4/432 | 0.9143 | 0.7105 |
| **zQ175** | 22.65 | 3.28 | 1.34 | 22.63 | 19.21-26.09 | 6/681 | | |

| **C) human muscle DAPI area (µm$^2$)** | | | | | | |
|---|---|---|---|---|---|---|
| | **mean** | **±SD** | **±SEM** | **median** | **95% CI** | **n/N** |
| **control** | 21.25 | 2.85 | 1.27 | 21.2 | 17.72-24.79 | 328/5 |
| **pre-HD** | 23.89 | 7.38 | 3.3 | 21.4 | 14.72-33.05 | 270/5 |
| **early HD** | 25.77 | 3.74 | 1.67 | 27.42 | 21.12-30.42 | 249/5 |
| ***Kruskal-Wallis test** p=0.3304 | | | | | | |
| **p values from Dunn's multiple comparison**<br>**control vs. pre-HD** >0,9999; **control vs. early HD** 0.4127; **pre-HD vs. early HD** 0.8665 | | | | | | |

| **E) % nuclei with nucleolar NPM1** | | | | | | | | |
|---|---|---|---|---|---|---|---|---|
| | **mean** | **±SD** | **±SEM** | **Median** | **95% CI** | **N/n** | ***MWU** | **t-test** |
| **control** | 19.72 | 8.20 | 3.35 | 21.91 | 11.12-28.33 | 6/672 | **0.0095** | 0.0101 |
| **TIF-IA$^{D1Cre}$** | 35.87 | 6.05 | 3.02 | 36.11 | 26.25-45.49 | 4/449 | | |

| **E) NPM1 area (µm$^2$)** | | | | | | | | |
|---|---|---|---|---|---|---|---|---|
| | **mean** | **±SD** | **±SEM** | **Median** | **95% CI** | **N/n** | ***MWU** | **t-test** |
| **control** | 2.50 | 0.53 | 0.24 | 2.50 | 1.84-3.17 | 5/131 | **0.0317** | 0.0171 |
| **TIF-IA$^{D1Cre}$** | 3.52 | 0.42 | 0.21 | 3.49 | 2.85-4.20 | 4/161 | | |

**Table 13**

Data and statistics for graphs shown in **Suppl. Figure 4F**, n: number of DAPI positive muscle nuclei, N: number of control individuals and Huntington's disease (HD) patients. * Statistical significance is shown according to the Kruskal-Wallis test.

| **F) % nuclei with nucleolar NPM1** | | | | | | |
|---|---|---|---|---|---|---|
| | **Mean** | **±SD** | **±SEM** | **median** | **95% CI** | **n/N** |
| **control** | 30.98 | 4.45 | 1.99 | 30.03 | 25.46-36.5 | 991/5 |
| **pre-HD** | 22.77 | 3.61 | 1.61 | 23.02 | 18.29-27.25 | 844/5 |
| **early HD** | 17.55 | 1.35 | 0.60 | 17.02 | 15.88-19.23 | 794/5 |
| **Kruskal-Wallis test p<0.0001** | | | | | | |
| **p values from Dunn's multiple comparison:**<br>**control vs. pre-HD** 0.2691**, control vs. early HD 0.0021, pre-HD vs. early HD** 0.2691 | | | | | | |

| **F) NPM1 area (µm$^2$)** | | | | | | |
|---|---|---|---|---|---|---|
| | **mean** | **±SD** | **±SEM** | **median** | **95% CI** | **n/N** |
| **control** | 3.59 | 0.68 | 0.30 | 3.77 | 2.75-4.43 | 991/5 |
| **pre-HD** | 2.79 | 0.81 | 0.36 | 2.52 | 1.78-3.79 | 844/5 |
| **early HD** | 2.25 | 0.23 | 0.10 | 2.23 | 1.97-2.54 | 794/5 |
| **Kruskal-Wallis test p=0.0236** | | | | | | |
| **p values from Dunn's multiple comparison:**<br>**control vs. pre-HD** 0.4719; **control vs. early HD 0.0267**; **pre-HD vs. early HD** 0.6880 | | | | | | |
